# A combined experimental-computational approach uncovers a role for the Golgi matrix protein Giantin in breast cancer progression

**DOI:** 10.1101/2022.04.25.489358

**Authors:** Salim Ghannoum, Damiano Fantini, Muhammad Zahoor, Veronika Reiterer, Santosh Phuyal, Waldir Leoncio Netto, Øystein Sørensen, Arvind Iyer, Debarka Sengupta, Lina Prasmickaite, Gunhild Mari Mælandsmo, Alvaro Köhn-Luque, Hesso Farhan

## Abstract

Few studies so far have investigated the impact of different cell migration traits on tumor progression. To address this, we developed a mathematical model wherein cells migrate in two-dimensional space, divide, die or intravasate into the vasculature. Exploring a wide range of speed and persistence combinations, we find that tumor growth positively correlates with increasing speed and higher persistence. As a biologically relevant example, we focused on Golgi fragmentation induced by depletion of Giantin, a Golgi matrix protein, the downregulation of which correlates with poor patient survival. Applying the migration and invasion traits of Giantin depleted cells to our mathematical model, we predict that loss of Giantin increases the number of intravasating cells. This prediction was validated, by showing that circulating tumor cells express significantly less Giantin than primary tumor cells. Altogether, our computational model identifies cell migration traits that regulate tumor progression and uncovers a role of Giantin in breast cancer progression.

## 1. Introduction

Breast cancer is the most commonly occurring cancer in women and the fifth most common cause of cancer-related death (1,2). The formation of metastases due to the spread of tumor cells to distant organs, is the major factor that determines mortality in breast cancer patients (3,4). Metastatic dissemination of breast cancer cells is a multi-step process requiring the acquisition of a motile phenotype, the ability to invade the surrounding tissue, break into vessels and eventually extravasate and colonize distant organs (5). Major research efforts in the past decades led to a firm understanding of the role of cell division, apoptosis and invasion in tumor progression and metastatic dissemination. However, there is surprisingly little knowledge of whether and how tumorigenesis is affected by different migration traits such as speed and directional persistence, *i*.*e*. the tendency to keep or change the direction of motion. Furthermore, it is not well-understood what the differential contribution of cell migration and invasiveness is to the ability of cancer cells to escape from the primary tumor site.

The Golgi apparatus (hereafter referred to as the Golgi) is a major cellular organelle that is involved in the regulation of membrane trafficking and post-translational modifications. In addition, the Golgi plays an important role as a regulator of directional cell migration (6-8). Under normal conditions, the Golgi is an intact single-copy organelle comprised of stacks of flattened cisternal membranes that are laterally linked to form the Golgi ribbon. This structure is maintained by Golgi-matrix proteins. Giantin is a such a Golgi-matrix protein that was shown to be important for maintaining the structural integrity of the Golgi (9-12). Several alterations in cultured cancer cells were reported to result in fragmentation of the Golgi (reviewed in (13)). Moreover, RNAi screening approaches have identified a wide range of conditions that affect the morphology of the Golgi (14-17). Golgi fragmentation has been reported to cause defects in cell polarization and therefore a delay in cell migration in wound closure assays (6,8,18). Our previous work indicates that that loss of a Golgi matrix protein GM130 in breast cancer cell lines is associated with altered migration and invasion (18). However, this analysis was so far only performed in cell lines and a confirmation of the pathophysiological significance of Golgi fragmentation in patient samples remains to be shown. Moreover, it does not answer the fundamental question of how cell migration affects tumor progression.

In the present study, we used *in silico* simulations to analyze the impact of a wide range of speed-persistence combinations of the shape and size of tumors and on intravasation of cancer cells. To this end, we constructed a mathematical model that accounts for migration, proliferation, invasion and death of cancer cells in a two-dimensional space. This model predicted that speed and persistence are positively correlated with the growth of the primary tumor. We then fed this model with experimental measurements of cell migration and invasion to study the impact of Giantin depletion on tumor progression. The model predicted that Giantin-depleted tumors, while smaller, are nevertheless capable of seeding more intravasating cells. This conclusion was then substantiated by showing that circulating tumor cells have lower Giantin levels than primary tumor cells. Our combination of modeling and experimental validation provides a possible explanation for the poor survival of breast cancer patients with low Giantin levels.

## 2. Materials and methods

### 2.1. Cell culture and transfection

MDA-MB-231 and BT549 were cultured in RPMI supplemented with 10% FCS and 1% penicillin/streptomycin (GIBCO). Cells were transduced with NucLight green lentivirus (Essen BioScience, 4624). Infections were carried out using a multiplicity of infection (MOI) of three transducing units per cell. The growth medium was changed after 24h of transduction and cells were split after 72h. The transduced cells were sorted via fluorescence-activated cell sorting (FACS) using a BD FACSAriaTM cell sorter. Sorted cells were tested for mycoplasma. For Giantin (GOLGB1) knockdown experiments, BT549 cells in optimal growth phase expressing GFP localizes to the nucleus were semireverse-transfected with siGENOME SMARTpool GOLGB1-siRNA (Dharmacon) with a final concentration of 15nM using HiPerfect (Qiagen, 301704) according to the manufacturer’s instructions. The SMARTpool of GOLGB1-siRNA contains 4 individual siRNAs with the following sequences; GAACUAGAGUCUCGGUAUA, UAAGAAUUGCAACCUAA, GUACACAGGUUAAGUGCUU and GAAGGUCUGUGAUACUCUA. The level of Giantin expression was determined by qRT-PCR. Total RNA was extracted using a GenElute^™^ Mammalian Total RNA Miniprep Kit (Merck, RTN70), and cDNA was reverse-transcribed using a High Capacity cDNA Reverse Transcription Kit (Thermo Fisher Scientific, Cat # 4368814). The expression of Giantin was determined using SensiFAST SYBR Hi-ROX (NordicBio, BIO-92002) and normalized to ACTIN using commercially available primers (Qiagen; for Giantin, QT00038087 and TBP, QT00000721).

### 2.2. Confocal immunofluorescence microscopy

GFP-BT549 cells were grown on glass cover-slips placed in a 6-well plate, and then fixed with 3% paraformaldehyde for 20 min at RT. Afterward, cells were washed in PBS with 20 mM glycine followed by incubation for 5 min at RT in permeabilization buffer containing PBS with 0.2% triton X100. Then, the fixed cells were incubated with mouse monoclonal anti-GM130 antibody (BD-Biosciences/Puls Medical, 610823) dilution 1:1000. After incubation with primary Abs at 37°C for 1 h followed by washing with PBS three times, cells were incubated for another 1h in goat anti-mouse secondary antibody, Alexa Fluro 488 (Thermo Fisher Scientific, A-11001) diluted in 3% BSA in PBS. Finally, cells were mounted in polyvinyl alcohol with DABCO antifade and imaged. Imaging was performed on Zeiss LSM700 laser scanning confocal microscope. All images were acquired randomly using a Plan-Aphochromat 63x Oil Ph3 M27 objective (NA 1.4).

### 2.3. Two-dimensional dense random migration assay

For optimal tracking efficiency we generated 3 populations; BT549 + GFP-BT549, BT549 + Giantin-silenced GFP-BT549, and MDA_MB_231 + GFP-MDA_MB_231. These cells were co-cultured at a ratio of 3:1 respectively in 96 well image-lock plates (EssenBio, 4739, Lot#17040501) for 24 hours at 37°C and 5% CO_2_. Then cells were scanned at ten-minute intervals over 24h in Essen BioScience’s IncuCyte S3 with objective lenses 10x (NA; 0.95; Image Resolution: 1.24um/pixel) using both the phase channel (HD Phase imaging in gray values) and green channel (Emission Wavelength: 524nm; Excitation Wavelength: 460nm; Exposure Time: 200ms). Images were collected using a Basler Ace 1920-155um camera with CMOS sensor. Cellular viability was assessed throughout the course of the scanning by comparing the phase cellular morphology between mother cells and their corresponding GFP cells. ImageJ was used to linearly correct the brightness and contrast of the images and merging them as multi-TIFF stack. All the images collected from the same experiment were modified the same way to leave the information unaltered. GFP cells were tracked and subjected to trajectory analysis using the in-house developed *cellmigRation* package as described in the next section (19)

### 2.4. The *cellmigRation* package

The *cellmigRation* is an open-soure R package that we developed to analyze cell movements using TIFF images captured over time. Its latest stable version is hosted on the peer-reviewed R/Bioconductor repository for easy installation and usage (DOI: 10.18129/B9.bioc.cellmigRation). The software includes two modules aimed at *i)* data import and pre-processing; and *ii)* advanced analytics and visualization. Image pre-processing (Module 1) is performed similar to what described by *DuChez B*, 2018 (20). Briefly, we imported the *FastTracks* software (https://www.mathworks.com/matlabcentral/fileexchange/60349-fasttracks) from *Matlab* to R and added functionality to automate some of the initial optimization steps (background correction, signal detection). Similar to *FastTracks*, our R library can import TIFF files, perform background removal, detect signal peaks corresponding to fluorescent particles in each stack of the image and record the corresponding centroid coordinates. Next, cell trajectories are computed by connecting the closest centroids from consecutive stacks provided that they are located within a user-defined maximum displacement radius. Cell tracking data can be used as input for the advanced analytic and deep trajectory analysis functions (Module 2). Our software can compute several metrics and statistics. Specifically, *cellmigRation* includes functions dedicated to the analysis of *i)* cell persistence and speed; *ii)* directionality; *iii)* mean squared displacement; *iv)* direction auto-correlation; and *v)* velocity auto-correlation. Module 2 also includes functions aimed at exporting data, automatically building cell-based and population-based visualizations and generating 3D interactive plots. Cell trajectory Principal Component Analysis (PCA) and Clustering are supported as well.

### 2.5. Gelatin degradation assay

Sterile cover-slips (12 mm, #1 Menzel™ Microscope Coverslips) were coated with a thin layer of 0.2 mg/ml Oregon Green 488-gelatin (Life Technologies, G13186) for 20 min, fixed in 0.5% glutaraldehyde for 40 min and washed three times with sterile PBS. Cover-slips were then transferred to a 24-well plate and incubated in complete growth medium for 30 min at 37 °C. BT549 cells transfected with either Ctrl-siRNA or GOLGB1-siRNA (Dharmacon) for 48h were seeded on top of the cover-slip at a density of 7.5×10^4^ cells/well and incubated overnight. Cells were then processed for anti-GM130 immunofluorescence staining. Images were randomly acquired on Andor Dragonfly spinning disk using a Nikon Ti2 inverted optical microscope equipped with a 60 × TIRF objective (Plan-APOCHROMAT 60 × /1.49 Oil). Fluorescence was collected using an EMCCD camera (iXon Ultra 888, Andor). To quantify the gelatin degraded areas per image, appropriate functions of ImageJ were used. The measured area was divided by the number of cells in the corresponding image. Kruskal-Wallis rank sum test was used to compare the gelatin degraded areas between the two conditions.

### 2.6. Patient-derived breast cancer xenografts

Patient-derived xenografts (PDX) were established at Institute for Cancer Research, Oslo University Hospital (MAS98.12 and MAS98.06) or the Institute Curie, France (HBCx-34 and HBCx-39) as previously described (21,22). Briefly, primary mammary tumor specimens were implanted into immunodeficient mice. After initial establishment, the tumor tissue was serially transplanted into mammary fat pads of nude athymic mice. Based on histopatological and molecular characterization, the tumors were classified as triple-negative basal like (MAS98.12 and HBCx-39) and hormone receptors positive luminal B breast cancer (MAS98.06 and HBCx-34) (21,23,24). our μm thick paraffin sections were rehydrated in a serial dilation of ethanol and boiled in TRIS-EDTA (pH 9.0) for antigen retrieval. The tissue sections were permeabilized with triton X-100 and then incubated at RT for 2 hours with anti-Giantin antibody (Nordic BioSite, bs-13356) dilution 1:1000. Then the tissue sections were washed with PBS three times, cells were incubated for another 1h in goat anti-mouse secondary antibody, Alexa Fluro 488 (Thermo Fisher Scientific, A-11001). The sections were incubated with Hoechst 33342 dye (Thermo Fisher Scientific, H3570) to stain the nuclei. The sections were rinsed with Milli-Q water and embedded in polyvinyl alcohol mounting medium (Sigma Aldrich, 10981). Fluorescence images were randomly acquired on on Andor Dragonfly spinning disk using a Nikon Ti2 inverted optical microscope equipped with a 60×TIRF objective (Plan-APOCHROMAT 60× /1.49 Oil). Fluorescence was collected using an EMCCD camera (iXon Ultra 888, Andor). To quantify the Golgi state, Golgi structure was assessed by visual inspection and categorized as intact, fragmented or undetermined.

### 2.7. Breast tissue array

Invasive ductal carcinoma and adjacent normal breast tissue samples were obtained from US Biomax, BC08118a. The slide was baked at 60 °C for 2h followed by incubation in Neo-clear (Sigma 109843) for 5min. The slide was re-hydrated in serial dilutions of Ethanol and then boiled in Tris/EDTA (pH 9,0) for 15min. After washing with PBS for 5min the slide was permeabilized using triton X-100 (0,25%) for 8min and then blocked in 5% BSA (in PBST) for 1h at RT. To stain the Golgi, the slide was incubated with anti-Giantin antibody (Nordic BioSite, bs-13356) in 1% BSA (PBST) for 2h at RT. After three times washing with PBST for 5min the slide was incubated for another 1h in goat anti-mouse secondary antibody, Alexa Fluro 488 (Thermo Fisher Scientific, A-11001). After several washing steps, the Hoechst 33342 dye (Thermo Fisher Scientific, H3570) was used to label the nuclei. Finally, the slide was rinsed with Milli-Q water and embedded in polyvinyl alcohol mounting medium (Sigma Aldrich, 10981).
 The fluorescence images were acquired and analyzed as previously described (section 2.6).

### 2.8. Circulating tumor cells/clusters (CTCs) datasets

To profile the expression of Giantin across normal and breast cancer cells in addition to circulating breast tumor cells, we used two single cell RNA-seq (scRNA-seq) datasets. The first dataset consists of normal and breast cancer cells. Data are available in the GEO database with accession numbers GSE114727 (25). The other dataset consists of single circulating tumor cells collected from the blood of breast cancer patients and obtained from several studies (26-33) compiled in our previous work (34). Data are available in the GEO database with accession numbers GSE51827, GSE55807, GSE67939, GSE75367, GSE109761, GSE111065, GSE86978 and PRJNA471754. After downloading raw scRNA-seq read count data we followed a standard pre-processing pipeline to filter the cells and genes as described in our previous work (34-36). After filtering the cells we performed the median by ratio normalization method followed by log transformation (36). For comparison purposes among circulating tumor cells, normal and breast cancer cells, cells with Giantin expression as zero were excluded from further downstream analysis.

### 2.9. Computational model of tumor progression

To investigate the impact of speed and directional persistence of breast cancer cells on tumor growth, we use a 2D stochastic cell-based model inspired by the persistence random walk model (37). Our newly developed model accounts for cell migration, proliferation, intravasation and death.

The model is based on 8 main assumptions: i) Cells are individual active agents that, for simplicity, live in a 2D section. ii) Cell movement is characterized by a given speed and directionality. iii) The 2D tissue section receives nourishment from functional blood vessels that are assumed to be perpendicular to the tissue section. iv) Cells in the nourished zone adjacent to a functional blood vessel are more likely to proliferate faster and move within this zone than leave it. v) Cell division depends on several factors, including proximity to blood vessels, local cell density and the time since the division of the corresponding mother cell. vi) Cells that spend the G1 phase in the nourished zones can have a shorter cell cycle length. vii) Cells become necrotic and die if they are not able to divide due to lack of space after several cell cycles. Senescent cells are also considered. Those are cells that are in a transition state between a proliferating state and a total cell cycle arrest. They can shift back to proliferating state if the local cell density changes. viii) Cells moving towards and eventually contact blood vessel can intravasate.

The aforementioned assumptions were translated into a stochastic mathematical and computational model that is fully described and parameterized in the Supplementary Material. Briefly, cells are described as circular agents of a given diameter and their position is determined by the coordinates of the center of the circle in a 2D computational domain. Cells cannot overlap but they can cross each other during migration. Cell movement is simulated according to speed profiles computed following an empirical distribution. The directionality of a cell is determined in term of persistence ratios, which range between 0 and 1. A persistence ratio of 1 refers to a very persistent cell, which maintained a preferred overall direction of movements within bilateral 90 degrees. The probability of a cell to divide depends on the local cell density, the time since the division of the original agent and the proximity to a blood vessel. We assumed that the proliferation rate is 30% higher in well-nourished spacious zones around cross-sections of blood vessels. This assumption was estimated empirically and is based on literature (38-40). The probability of undergoing cell division increases significantly when the optimal cell-cycle time is achieved. Additionally, cells are removed from the field to simulate i) cell death if a cell is not undergoing cell division after a number of cell-cycles because of lack of available space. ii) intravasation if a cell succeeded penetrating into the lumen. In our model, the ability of a cell to move, divide, intravasate and leave a well-nourished zone is a stochastic event. Such stochasticity is achieved by generating a uniform random number between 0 and 1 per cell per process per iteration. A process is triggered when the random number is equal to or less than the corresponding parameter value, representing the probability of the event to take place. Periodic boundary conditions were imposed to avoid losing cells going out of the simulation field.

In each simulation, which is characterized by certain speed and directional persistence, we accounted for the following outcomes: the number and distribution of cells in the 2D domain, the number of intravasating cells, the proliferation outcome and the mean time spent in the nourished zones. In this way we were able to estimate the optimal speed-persistence combination that leads to bigger tumors and more intravasation. We also used the model to study the pathological relevance of Golgi fragmentation in breast cancer. All the modeling simulations were implemented using the high-performance computing at University of Oslo (Abel computer cluster). The source code to simulate the model is available at https://github.com/ocbe-uio/Cancer_simulator.

## Results

### The effect of cell speed and persistence on tumor progression

Two of the most intuitive parameters to characterize cell movement are the speed and directional persistence. We first investigated the impact of these two parameters on tumor growth and intravasation using computational modeling. Our model accounted for cell migration, proliferation, intravasation, and cell death (see the Methods section and Supplementary Material for a full description of the model and its parameterization). Briefly, cells were freely moving in a two-dimensional space, were assigned a certain rate of division, and were assumed to die if they had not performed a division for longer than three doubling times. This allowed us to simulate cell death and thereby tumor necrosis in poorly nourished areas. To account for growth advantage and attraction of the well-nourished areas close to blood vessels, cells were assigned a lower likelihood to leave this area and were given a 30% growth advantage (faster doubling). We implemented extensive numerical simulations of 130 speed-persistence combinations. We used live epifluorescence imaging of cells (BT549 and MDA-MB-231) migrating freely in 2D followed by automated tracking to infer a realistic range of parameter space for persistence and speed (see the supplementary materials for the experimental outcome). For each simulation of a 90 days period, we computed the changes in the number and distribution of cancer cells. We characterized the spatial cell distribution using the uniformity index (UI) (41-43). High UI indicates a homogeneous distribution all over the simulation field, whereas low UI is an indication for cluster formation. Due to the model stochasticity, simulations were repeated ten times for each speed-persistence combination (see supplementary video 1).

Simulations showed that cell speed and persistence have a profound effect on the distribution of cancer cells generating tumors with different uniformity indices (UIs) (Fig. 1A). Our model also recapitulated tumor necrosis in the center of compact tumors with a high UI (see inset in Figure 1A and Fig. S1A). A key prediction of our in silico simulations is that the size of the tumor increases with both speed and persistence of cell migration (Supplementary Fig. S1C). The impact of speed was stronger than that of persistence. When a fixed persistence value was used, increasing the speed had a major impact on the number of cells in the tumor (Fig. 1B). The impact of persistence on the size of the tumor was barely noticeably at low speeds (Supplementary Fig. S1D).

**Fig. 1.**
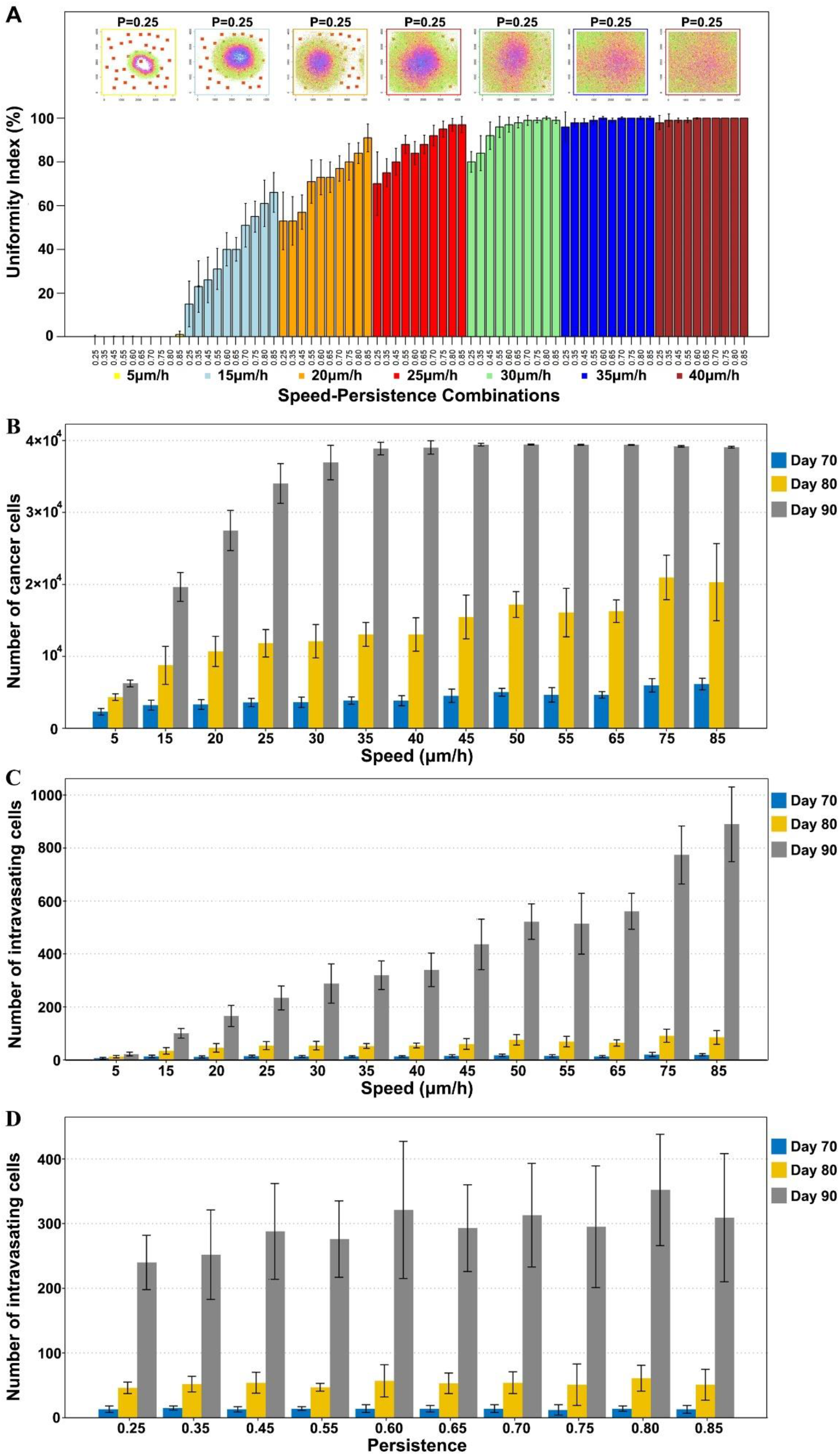
Effect of cell speed and persistence on tumor growth and intravasation. (A) Upper plots illustrate how the distribution of cancer cells at the end of the simulation changes for different speeds and the same fixed persistence (P = 0.25). Bars represent mean uniformity index of ten simulations. Different colors refer to different speeds. Within one color, persistence gradually increases. (B) Number of cancer cells at different time points across gradual speed but fixed persistence (P = 0.45). (C) Number of intravasating cells at different time points across gradual speed but fixed persistence (P = 0.45). (D) Number of intravasating cells at different time points across gradual persistence value but fixed speed of 30um/h. Error bars represent the standard deviation of the data.

Cells moving towards and contacting a blood vessel were assigned a certain probability of leaving the tumor by intravasation. Simulations showed that the number of intravasating cells increases with speed only (Supplementary Fig. S1E). Under the same persistence condition, speed had a strong impact on the number of intravasating cells, but no effect on persistence was observed when the speed was kept constant (Fig. 1C&D). This is probably because speed has a stronger impact of the total number of cells in the tumor and it is intuitive to assume that larger tumors will produce more intravasating cells.

### Golgi fragmentation in breast cancer tissue

Next, we sought to explore the biological significance of our computational model. As stated above, alterations of Golgi structure are well known to affect cell migration (6,15). Thus, we decided to test pathophysiological relevance of our model predictions by investigating cell migration traits associated with a condition that induces Golgi fragmentation. Although the role of the Golgi in tumor progression has been discussed in the literature (6,13,44,45), there is surprisingly little experimental evidence for structural alterations of the Golgi in patient-derived tumor tissue. Golgi fragmentation has been reported in colorectal cancer (46) and prostate cancer (47), but to the best of our knowledge no analysis of a Golgi structure in breast cancer tissue is available. Thus, before directly investigating a specific condition that induces Golgi fragmentation, we wanted to assess Golgi structure in patient-derived xenograft (PDX) samples from triple-negative basal-like, or hormone receptors positive luminal B tumor tissue (Fig. 2A&B). In the triple-negative tumors, Golgi fragmentation was observed in 50% of the examined cells whereas only 41% of the hormone receptors positive tumors contained fragmented Golgi. The prevalence of this phenotype tempts to speculate about a relevance of Golgi fragmentation in breast cancer pathology. This notion is further supported by the observation of the higher prevalence of Golgi fragmentation in samples from invasive ductal carcinoma compared to adjacent normal breast tissue (Supplementary Fig. S2).

**Fig. 2.**
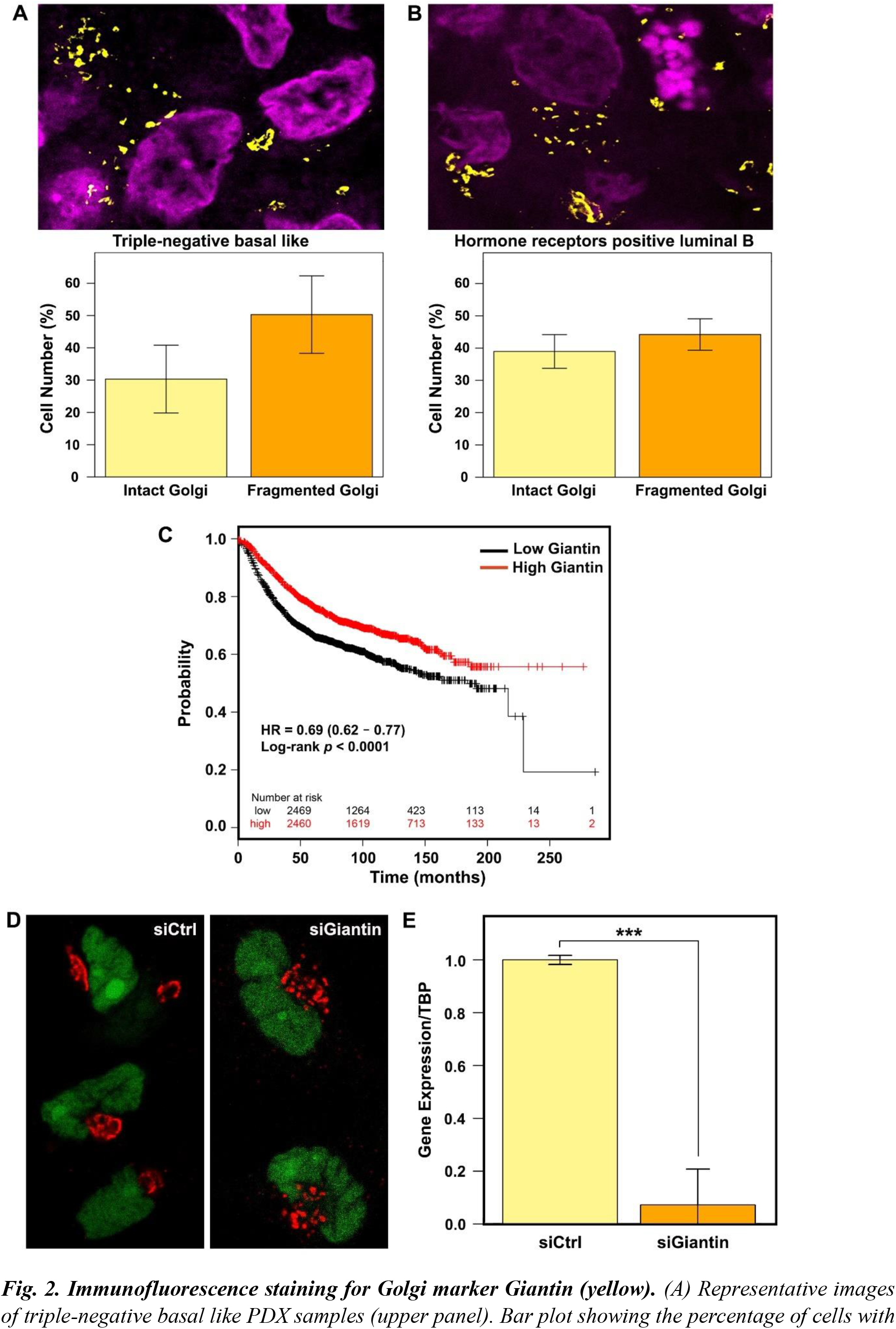
Immunofluorescence staining for Golgi marker Giantin (yellow). (A) Representative images of triple-negative basal like PDX samples (upper panel). Bar plot showing the percentage of cells with intact and fragmented Golgi (lower panel) in different triple-negative basal like PDX samples (MAS98.12 and HBCx-39; n = 4 with 25 fields each). (B) Representative images of hormone receptors positive luminal B PDX samples (upper panel). Bar plot showing the percentage of cells with intact and fragmented Golgi (lower panel) in different hormone receptors positive luminal B PDX samples (MAS98.06 and HBCx-34; n = 4 with 25 fields each). Error bars represent standard deviation of the data. (C) Kaplan–Meier survival curves in patients with high and low Giantin expression. The graph was generated using the kmplot database (https://kmplot.com/) (D) Representative image of BT549 cells stably expressing nuclear GFP were transfected with Ctrl or Giantin siRNA. After 72 h, cells were fixed and immunostained against GM130 to label the Golgi. (E) Barplot showing the average values of the relative gene expression of Giantin in 3 biological replicates. Standard deviation is reported above each bar. Gene expression level was quantified relative to reference gene TBP by ΔΔCq method. Statistical significance was determined using Kruskal-Wallis rank sum test (*** p < 0.001).

### Giantin is a Golgi protein that might be involved in breast cancer progression

Previous work has uncovered a wide range of genes that when depleted lead to Golgi fragmentation (14-16). However, because these data are from siRNA screens, it remains largely unclear whether these gene depletions induce Golgi fragmentation in a direct manner. Thus, we decided to focus on Golgi matrix proteins and tested whether their levels are linked to survival of breast cancer patients using the kmplot database (48,49). We focused on six matrix proteins that have a relatively well understood role in maintaining Golgi morphology (GOLGA1, GOLGA2, GOLGA3, GRASP55, GOLGB1 and GORASP1). With the exception of GRASP55 and GOLGA3, there was a significant trend indicating that low expression correlates with poor survival (Supplementary Fig. S3). This trend was very strong for Giantin (also known as GOLGB1, Fig. 2C) and therefore we decided to focus on this Golgi matrix protein for further studies. Silencing Giantin expression in BT549 cells induced a fragmentation of the Golgi (Fig. 2D&E and Supplementary Fig. S4A). To investigate the migratory pattern of BT549 cells upon Giantin knockdown, we analyzed the trajectories of cells that were freely migrating on a 2D surface. Cells were scaled and projected based on their root mean square speed and persistence ratio. Scaling was done by normalizing them to the mean values, which were assigned a value of 1. Using this approach, we defined four groups with distinct migration and persistence values: group 1 (low speed, low persistence), group 2 (low speed, high persistence). group 3 (high speed, low persistence) and group 4 (high speed, high persistence) (Fig. 3A). In comparison to control cells, BT549 cells lacking Giantin were preferentially found in the low-speed groups 1&2 (Fig. 3B,C&D; Supplementary Fig. S4B). On the other hand, cell persistence was not appreciably affected by the depletion of Giantin (Supplementary Fig. S4C&D). Seeing these experimental results through the lens of our computational model, the migration pattern of Giantin depleted cells would result in a smaller tumor, which we expect to seed less intravasating cells. This is difficult to reconcile with the fact that patients with lower Giantin exhibit lower disease-free survival rates (Fig. 2C). Our simulations were all carried out with a fixed rate of intravasation and therefore, it appears intuitive that smaller tumors would produce less intravasating cells. Thus, we asked to what extent must the rate of intravasation of a smaller tumor increase to produce more intravasating cells than a bigger tumor? We therefore re-run our simulations with adjustments of the rates of intravasation. To better link these new simulations to our experimental data, we took into account the number of cells with Golgi fragmentation in conditions of Giantin knockdown, where we observed that 65% of cells display a fragmented Golgi. We assumed that the cells with fragmented Golgi are the ones with the highest level of Giantin knockdown where the biological effects (*i*.*e*. slower cell migration) is also expected to be strongest. In the simulations, tumors with low Giantin, were considered to have 65% of cells with slower cell migration. Because control conditions exhibited 14% of cells with Golgi fragmentation, we used this value for the simulation of tumors with high levels of Giantin. The higher the number of cells with Golgi fragmentation, the smaller the tumor and the less intravasated cells it produced. This is to be expected, because Golgi fragmentation (induced by Giantin depletion) slows cells migration, which in our model reduces the size of the tumor, thereby having less cells than can intravasate. When we assigned the Giantin-depleted tumor a 2-fold higher intravasation rate, we observed that it produced a similar amount of intravasated cells compared to control conditions with only 14% fragmented Golgi (Fig. 3E). Of note, the tumor with 65% Golgi fragmentation was still smaller (Fig. 3F) indicating that the increase in intravasation rate can overcome the effect of the size of the primary tumor. Increasing the intravasation rate by 4-fold allowed the tumors with high degree of Golgi fragmentation to seed more intravasated cells (Fig. 3E), which again could not be explained by the tumor size (Fig. 3F). Based on this, we hypothesized that increasing the invasion rate by at least 4-fold allows smaller tumors to seed more circulating tumors cells, which might give rise to more metastasis and thus might explain poor patient survival.

**Fig. 3.**
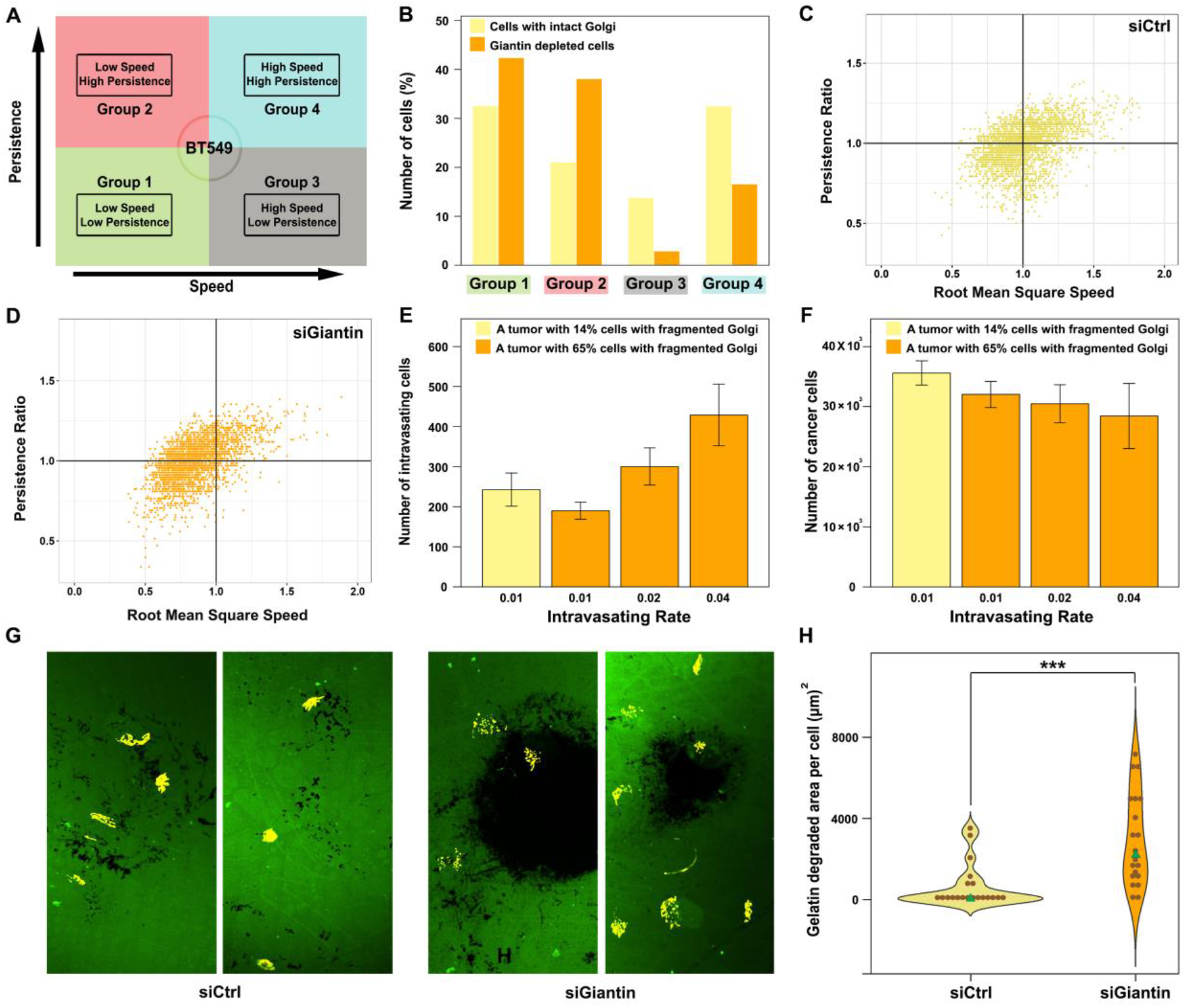
BT549 migration and invasion. (A) A schema showing the four groups of BT549 cells based on speed and persistence. (B) Bar plot showing the distribution of cells in each group. (C) The four migratory fashion groups of BT549 control cells. (D) The four migratory fashion groups of BT549 cells with Giantin knockdown. (F) The impact of increasing the intravasation rate of cells with fragmented Golgi on tumor growth. (G) Representative confocal immunofluorescence images of fluorescent gelatin degradation assay in BT549 cells. Degraded areas are visualized in black. The Golgi is visualized in yellow. (H) Violin plot showing the quantification of the gelatin-degraded area. Green tringles represent the median value. Black dots represent the gelatin-degraded area normalized to cell number. Statistical significance is assessed by Kruskal-Wallis rank sum test (*** p < 0.001), n= 20 microscopic fields from 2 independent experiments.

To experimentally test whether Giantin depleted cells exhibited higher invasion rates, we tested their ability to degrade extracellular matrix. To this end, cells were seeded on glass cover slips coated with fluorescent gelatin. By analyzing the pattern and size and the area of the digested gelatin we found that silencing Giantin resulted in a marked increase of the median ability of cells to degrade gelatin. This increase ranged from a few percent to more than 30-fold (see violin plot and representative images in Fig. 3G&H).

To predict the pathological relevance of Giantin loss in breast cancer we expanded our *in silico* model to account for a higher rate of invasion which we observed experimentally. The revised model had three additional assumptions: 1) each cell in the tumor has a probability of developing the low Giantin phenotype; 2) cells with low Giantin have higher invasiveness, which yields to higher intravasating capability; 3) cells follow the migratory phenotype of one of the four aforementioned groups based on the percent abundance of the groups. The *in silico* outcome of tumor progression in two independent populations (Supplementary video 2) that diverge in the percent abundance of cells with low Giantin (14% and 65%) shows significant difference in terms of uniformity index, number of generated cells and number of intravasating cells (Fig 4A). Furthermore, we tested the impact of a serial percent abundance values of cells with low Giantin (0%, 25%, 50%, 75% and 100%) on tumor progression (Supplementary video 3). The results show that the higher the abundance of cells with low Giantin, the lower the uniformity index and the smaller the tumor but with higher intravasation capability (Fig 4B-4D). The number of intravasating cells in a population of a thousand cancer cells was found to be statistically significant among all the different groups with the highest number (30 ± 4) in a pure population of cells with low Giantin expression (Supplementary Fig. S5). Thus, the combination of our mathematical and cell biological experiments provides a possible explanation for the poor disease-free survival of patients with low Giantin levels.

**Fig. 4.**
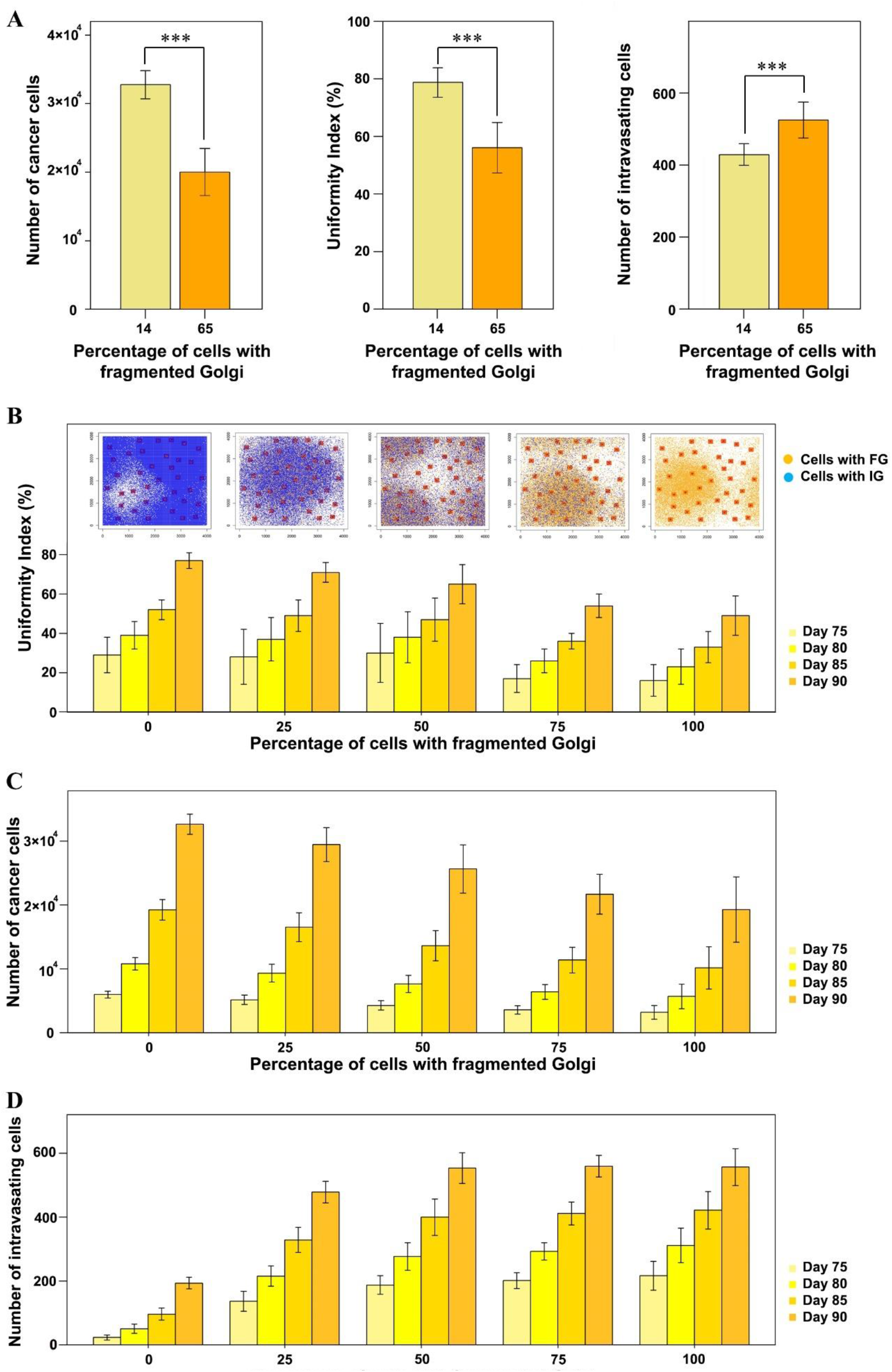
Impact of loss of Giantin on tumor growth and intravasation. (A) Modeling outcome of a tumor with 14% cells with fragmented Golgi versus a tumor with 65% cells with fragmented Golgi. Statistical significance was determined using two sample t-test (*** p < 0.001), (B) Upper plots demonstrate the distribution of cancer cells after 90 days from the tumor initiation with different percent abundance values of cells with fragmented Golgi. Cells with an intact Golgi (IG) are depicted in blue and cell with a fragmented Golgi (FG) are depicted in yellow. Bars represent mean uniformity index of ten replicates. Different colors refer to different time points. (C) Number of cancer cells at different time points across gradual percent abundance values of cells with fragmented Golgi. (D) Number of intravasating cells at different time points across gradual percent abundance values of cells with fragmented Golgi. Error bars represent the standard deviation of ten replicates.

### *Giantin* expression in breast cancer patients

Because our simulation results suggest that loss of Giantin increases the invasive capability of tumor cells to intravasate into the circulation, we would expect circulating tumor cells to have lower Giantin expression than the primary tumor. To test this hypothesis, first we investigated the expression of Giantin in circulating breast tumor cells (*n* = 1448) compared to a housekeeping gene (*ACTB*) (Fig. 5A). Then, we profiled the expression of Giantin in individual cancer cells (n = 168) and adjacent normal breast cells (*n* = 1500) collected from patients with breast cancer in addition to circulating breast tumor cells (*n* = 627). The single-cell RNA-seq analysis showed significant gradual decrement in mRNA levels of Giantin across the three datasets with lowest expression in the circulating breast tumor cells (0.24 ± 0.81) compared to primary breast cancer cells (0.99 ± 0.96) and their adjacent normal breast cells (1.84 ± 0.76) (Fig. 5B). On the other hand, we observed a marginal difference in the expression of a housekeeping gene (*ACTB*) (Fig. 5C).

**Fig. 5.**
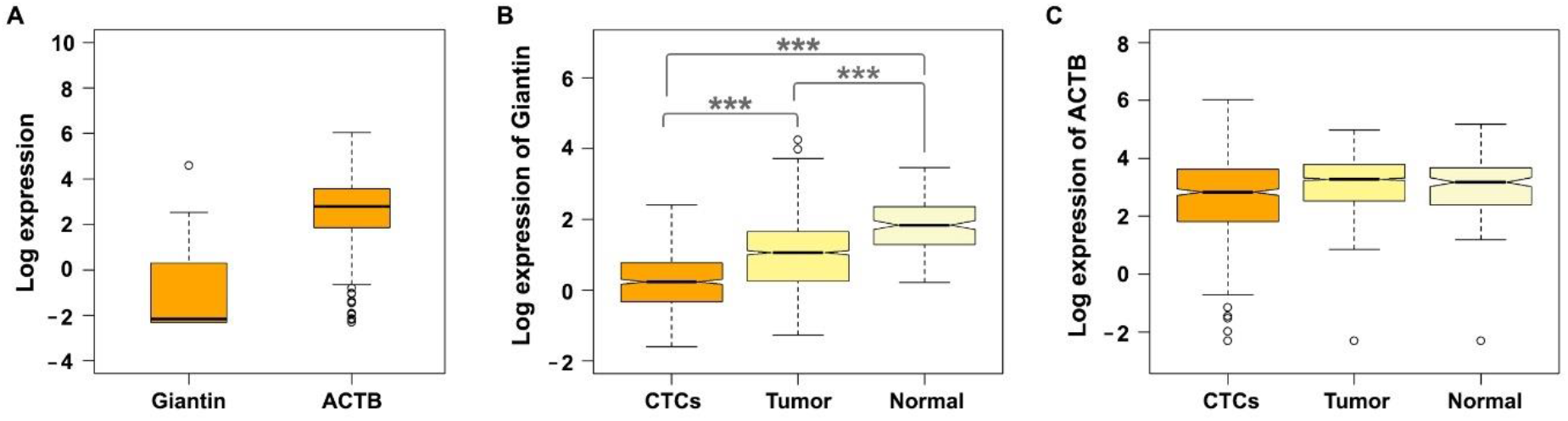
Giantin expression profile. (A) Boxplot showing the expression of Giantin and actin (ACTB) at single cell level in CTCs (n=1448). (B) Boxplot showing the expression of Giantin at single cell level in CTCs (n=627), breast tumor (n = 168) and adjacent normal breast (n = 1500). Statistical significance was determined using the Kruskal–Wallis rank sum test, followed by pairwise comparisons using the Wilcoxon rank sum test (*** p < 0.001). (C) Boxplot showing the expression of actin (ACTB) at single cell level in CTCs (n=627), breast tumor (n = 168) and adjacent normal breast (n = 1500).

## Discussion

In addition to state-of-the art experiments, mathematical and computational models provide insights into complex cancer processes including tumor growth (50,51), metastasis (52-54), treatment response (52,55-60) and cell migration and invasion in 2D and 3D matrices (61-64). In particular, cell-based models have been recently applied to investigate the roles of proliferation, migration, and necrosis in tumor progression of different cancers (reviewed in (65)). The present study represents a new step in understanding the impact of different traits of cell migration on the progression of breast cancer through an integrated experimental-computational approach. Thereby, we provide a possible explanation for the poor survival of breast cancer patients with low expression of Giantin. Loss of this Golgi matrix protein results in fragmentation of the Golgi apparatus, a phenomenon that we found to be very prevalent in patient-derived breast cancer specimens. Our computational approach shows that a simple model accounting for cellular activities and migratory characteristics can be used to study tumor size and the intravasating rate over a given time. Importantly, our model suggests that cells with lower speed lead to a smaller and more compact tumor. Because silencing Giantin led to a reduction in migration speed, our model predicts that tumors with low Giantin expression would be smaller in size. Thus, our computational model indicated the need for further experimental results to explain the clinical survival data. We then noted that silencing Giantin enhanced invasiveness of breast cancer cells and through further computational simulations, we found that this enhanced invasiveness could overcome the effect of the smaller size such that Giantin-low tumors would produce more circulating tumor cells. This notion is the observation that CTCs express lower Giantin levels than the primary tumor.

Several questions arise from our work. One question is whether the compactness of tumors with lower Giantin levels can be validated clinically. Based on this, it will be interesting to evaluate the chemosensitivity of tumors with different levels of compactness. It is possible that chemotherapeutics might have less access to compact tumors, thus making these tumors more likely to relapse after an initial round of therapy. Our mathematical model focused mainly on the cancer cells and did not consider the role of cancer associated fibroblasts or the tumor stroma. It is possible that Giantin-low tumors might be smaller, but also less compact due to the higher invasiveness. Indeed, less compact tumors were found to be associated with lymphovascular space invasion and the formation of lymph node metastases (66). Our model and the experimental follow-up do not allow to determine how the Giantin-low trait is being selected for. While the acquisition of a more invasive phenotype is an advantage for intravasation, it appears counterintuitive to select for a trait that would cause a growth disadvantage. However, it is possible that this switch between a pro-invasive trait and a pro-growth trait is a reflection of this phenotypic plasticity that is well known in tumors. It is also possible that the invasive phenotype might alter the interaction of cancer cells with their microenvironment and thereby alter tumorigenesis in a way that can only be investigated using in vivo imaging techniques and is therefore beyond the scope of the current study.

The exact explanation of the elevated invasion capability of cells with low Giantin is not yet clear but a few possible scenarios can be hypothesized. One possible explanation is that Giantin depleted cells exhibit a reduction in their speed, thus having more time for efficient degradation of the collagen while moving slowly on top of it. Another scenario is that silencing Giantin induces the expression of matrix metalloproteinases (MMPs) to degrade many components of the extracellular matrix. Finally, there is evidence that fragmentation of the Golgi enhances the rate of intra-Golgi trafficking (67). Although this has so far only been shown in the context of neurodegeneration, it is possible that a fragmented Golgi would accelerate the secretory transport of MMPs and thereby enhance extracellular matrix degradation.

Taken together, our work used a combination of mathematical modeling and wet-lab experiments and proposed that the speed of cancer cell migration has a major impact on the shape and size of a tumor and this may explain how loss of Giantin could contribute to the process of tumorigenesis.

## Acknowledgement

HF is supported by funding form the Norwegian Cancer Society (grants 182815 & 208015), by the Norwegian Research Council (grant 302452), by the Anders Jahre Foundation, and by the Rake og Otto Kr. Bruun legat. A.K.L acknowledge support by the center for research-based-innovation BigInsight funded by the Research Council of Norway under project number 237718. A.K.L and S.G were supported by the UiO:Life Science convergent environment PerCaThe. This work was performed on the Abel computer cluster, owned by the University of Oslo, and operated by the Department for Research Computing at USIT, the University of Oslo IT-department. http://www.hpc.uio.no/

## Notes

The authors declare no potential conflicts of interest

### Competing Interest Statement

The authors have declared no competing interest.

