## Supplementary Materials for "A combined experimental-computational approach uncovers a role for the Golgi matrix protein Giantin in breast cancer progression"

[1] Institute of Basic Medical Sciences, Department of Molecular Medicine, University of Oslo, Norway.
[2] Department of Urology, Northwestern University, Chicago, USA.
[3] Institute of Pathophysiology, Medical University of Innsbruck, Austria
[4] Oslo Centre for Biostatistics and Epidemiology, Faculty of Medicine, University of Oslo, Norway.
[5] Center for Lifespan Changes in Brain and Cognition, University of Oslo, Norway.
[6] Department of Computational Biology, University of Lausanne (UNIL), Lausanne 1015, Switzerland.
[7] Department of Computational Biology, Indraprastha Institute of Information Technology, New Delhi 110020, India.
[8] Department of Tumor Biology, Institute for Cancer Research, Oslo University Hospital, The Norwegian Radium Hospital, Oslo, Norway.
[ŧ] Shared first authorship
[*] Correspondence to:




**Supplementary Fig. S1.**

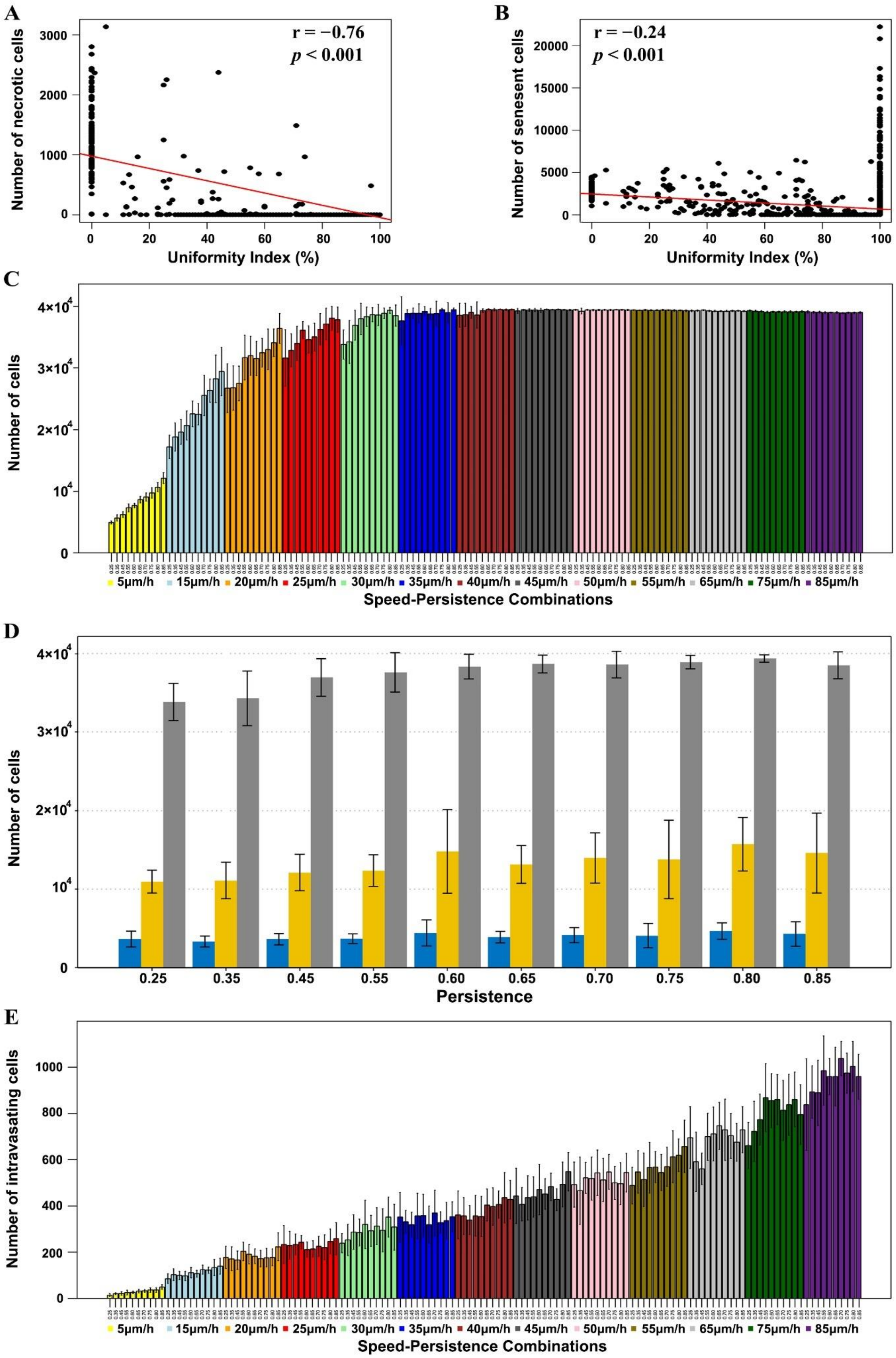

***Supplementary Fig S1. Impact of cell speed and persistence on tumor progression.*** *(A) Strong negative correlation between number of necrotic cells and uniformity index (Pearson correlation coefficient = -0.76). (B) Weak negative correlation between number of senescent cells and uniformity index (Pearson correlation coefficient = -0.24). (C) Number of cancer cells after 90 days from the tumor initiation across different speed-persistence combinations. (D) Number of cancer cells at different time points (blue: day 70, yellow: day 80 and gray: day 90) across gradual persistence value but fixed speed of 30um/h. (E) Number of intravasating cells after 90 days from the tumor initiation across different speed-persistence combinations. E*rror bars represent *the* standard deviation *of ten replicates.*

**Supplementary Fig. S2.**

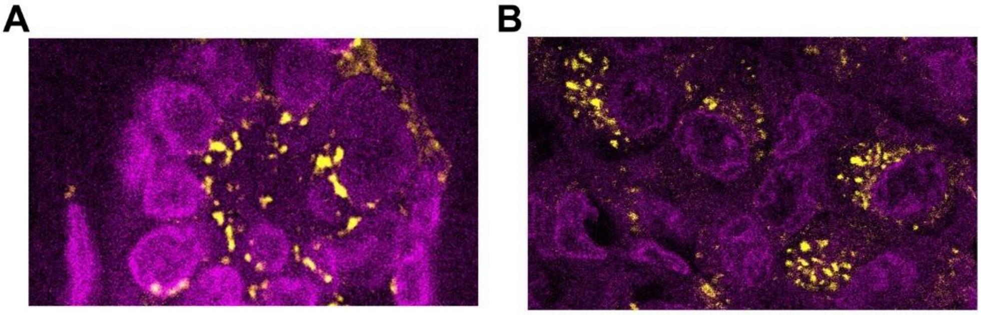


***Supplementary Fig. S2. Immunofluorescence staining for Golgi marker giantin (yellow).*** *(A) Representative image of adjacent normal breast tissue patient samples. (B) Representative image of invasive ductal carcinoma patient section.*

**Supplementary Fig. S3.**

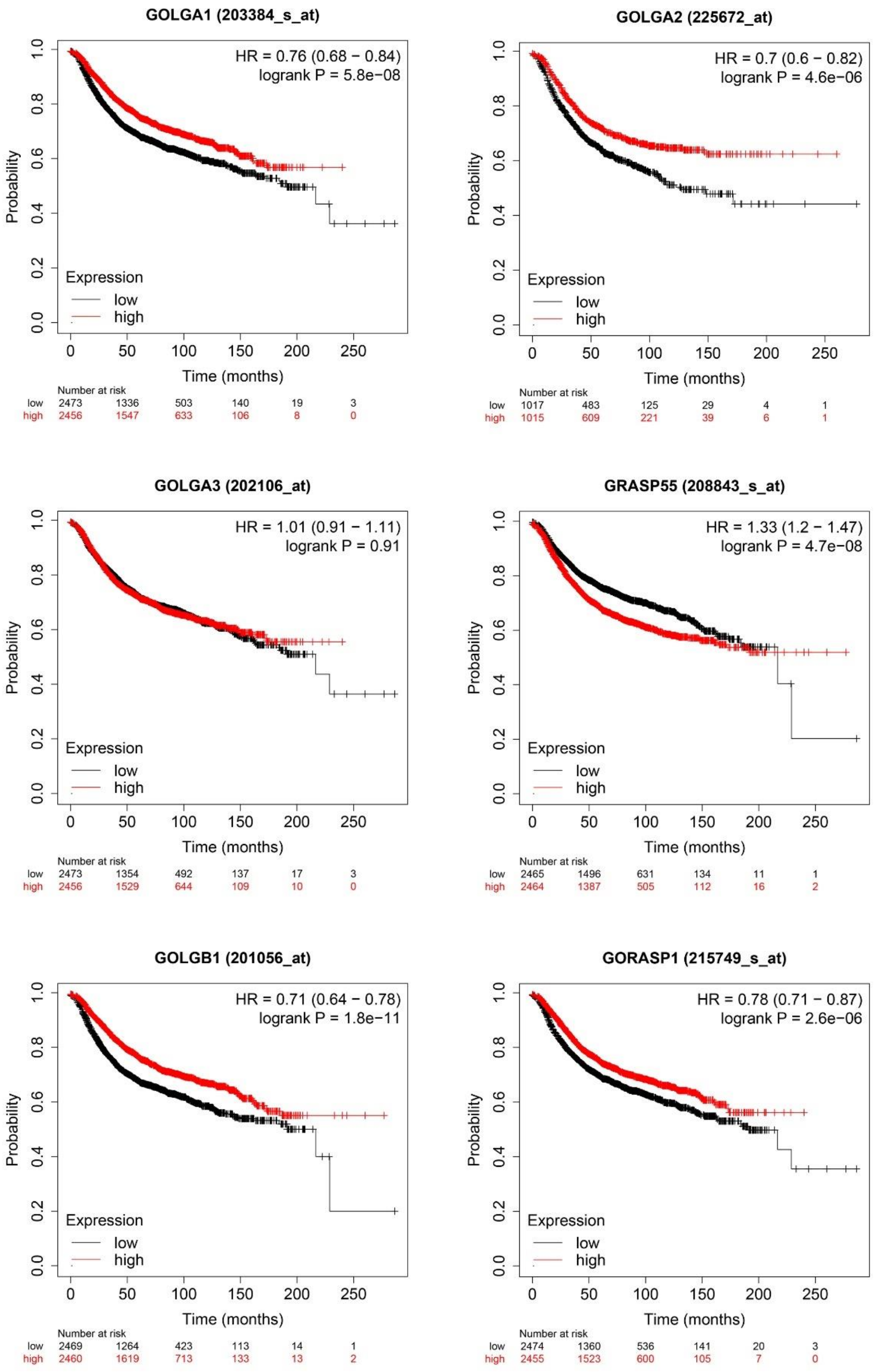


***Supplementary Fig. S3. Kaplan–Meier survival curves** for* six matrix proteins that have a relatively well understood role in maintaining Golgi morphology (GOLGA1, GOLGA2, GOLGA3, GRASP55, GOLGB1 and GORASP1).

**Supplementary Fig. S4.**

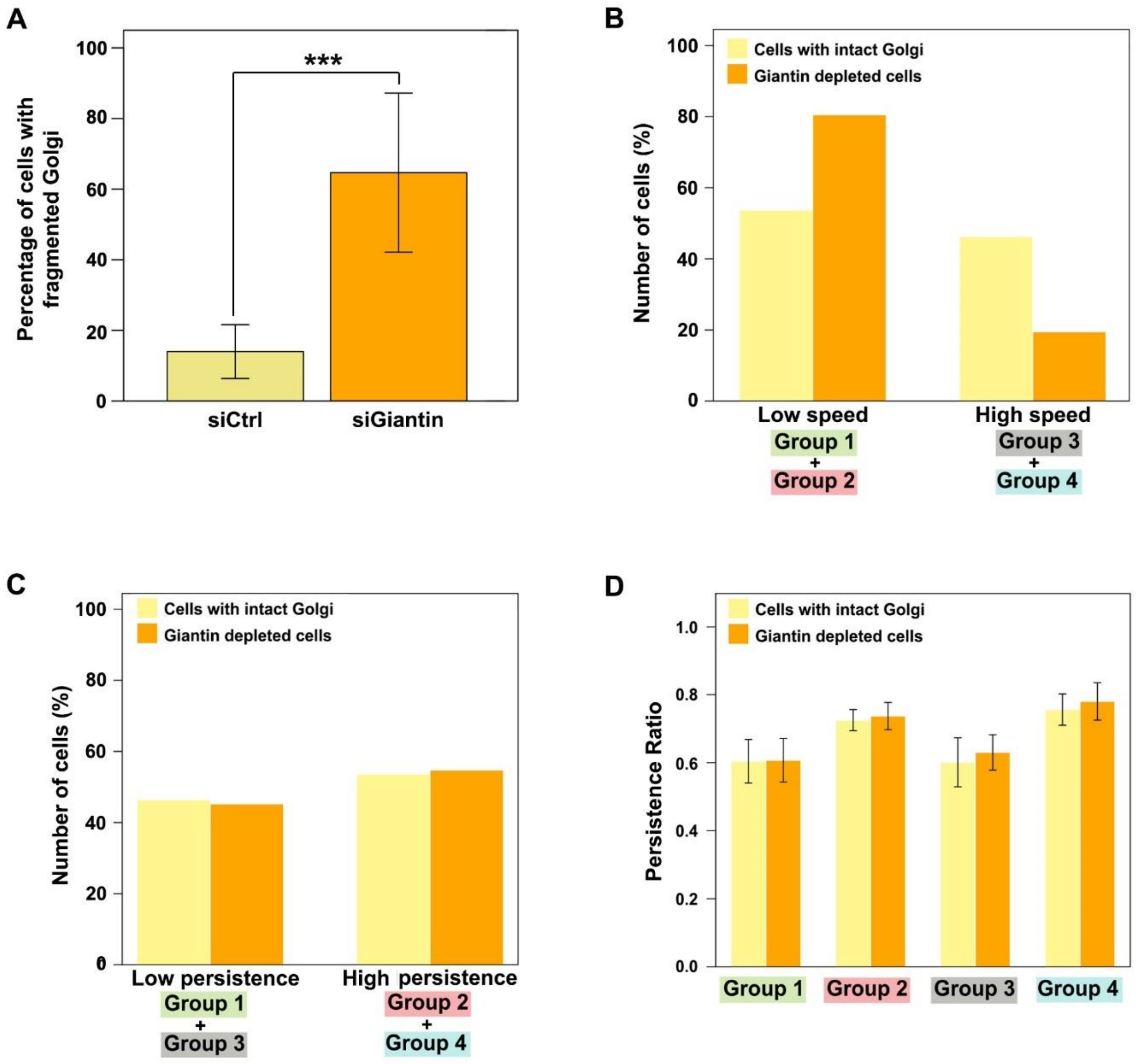


***Supplementary Fig. S4. Impact of Golgi fragmentation on cell migration.*** *(A) Percentage of cells denoted with fragmented Golgi in BT549 cells. (B) Bar plot showing the percentage of cells based on speed, low speed includes group 1&2 whereas high speed includes group 3&4 . (C) Bar plot showing the percentage of cells based on persistence, low persistence includes group 1&3 whereas high persistence includes group 2&4. (D) Bar plot showing the averaged persistence ratio per group*

**Supplementary Fig. S5.**

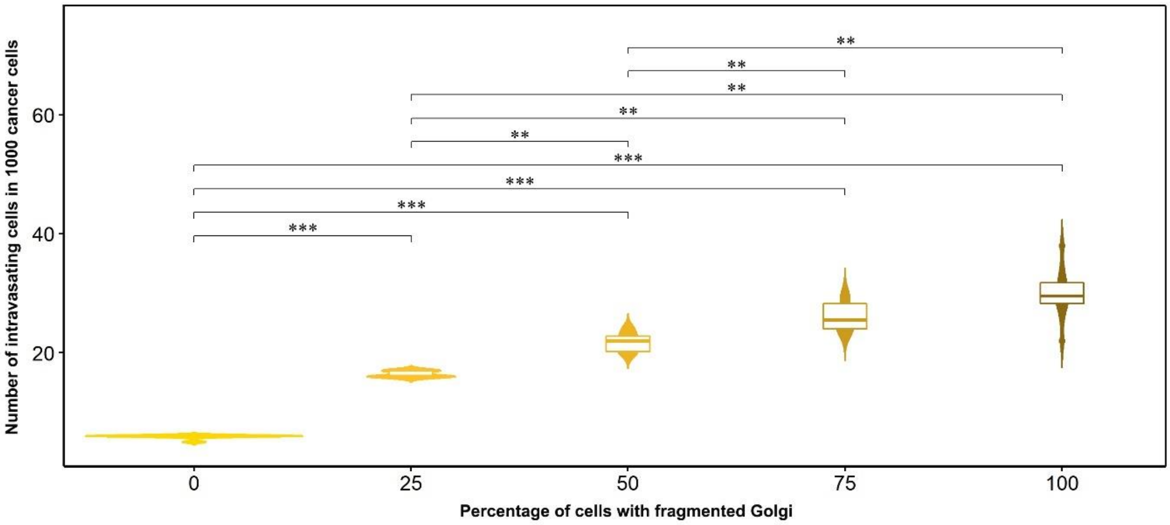


***Supplementary Fig. S5. Number of intravasating cells in a population of a thousand cancer cells.*** *Statistical significance was determined using the Kruskal–Wallis rank sum test, followed by pairwise comparisons using the Wilcoxon rank sum test with* Bonferroni Correction *(** $p < 0.01$, *** $p < 0.001$).*

### Computational model of breast cancer growth and progression

We use a stochastic cell-based model, built on the persistence random walk paradigm, to describe not only cell migration but also proliferation, intravasation, senescence and cell death. This section describes in full detail: 1) the model assumptions; 2) how those assumptions are translated into a mathematical and computational framework; 3) the used model parameters and initial conditions; and 4) all computational details needed to simulate the model.

#### 1. Model assumptions

Our stochastic model is based on 8 main assumptions:

i) Cells are individual active agents that, for simplicity, live in a 2D section.

ii) Cell movement is characterized by a given speed and directionality.

iii) The 2D tissue section receives nourishment from functional blood vessels that are assumed to be perpendicular to the tissue section.

iv) Cells in the nourished zone adjacent to a functional blood vessel are more likely to proliferate faster and move within this zone than leave it.

v) Besides proximity to blood vessels, cell divisions are also influenced by the local cell density and the time since the division of the corresponding mother cell.

vi) Cells that spend the G1 phase in the nourished zones can have a shorter cell cycle length.

vii) Cells become necrotic and die if they are not able to divide due to lack of space after a number of cell cycles. Senescent cells are also considered. Those are cell that are in a transition state between a proliferating state and a total cell cycle arrest. They can shift back to proliferating state if the local cell density changes.

viii) Cells moving towards blood vessel can intravasate.

#### 2. Model description

Here we translate the model assumptions into mathematical and computational framework.

i) Single cells are described as circular discrete agents with certain diameter (d). No two agents can occupy the same location simultaneously, but they can cross each other during migration. The position of each agent is determined by the coordinates of its center that can be at any point in a continuous 2D simulation field (F).

ii) Agents change their coordinates across discrete time steps by a given speed and directionality. Instead of having constant speed, the agents follow speed distributions computed empirically (Fig. S6A). Cells' directionality is determined in term of persistence ratio ($P_r$) which is computed as:

$$P_r = N_p/(N_p + N_d) \qquad \text{(Equation S1)}$$

where, $N_p$ is the number of persistent steps and $N_d$ is the number of deviating steps (Fig. S6B). Time-step movement vectors, referred to as steps, are classified either persistent or deviating based on the angular degree between the extension of the previous movement distance and the current one (Fig. S6B). Persistent steps are associated with acute angles ($< \pi/2$) whereas deviating steps are associated with obtuse angles ($\geq \pi/2$).

iii) For simplicity, cross sections of blood vessels ($L_i$, i=1…Nv), referred to as lumina, are stochastically distributed in the simulation field (F) and modelled by squares with fixed area (L).

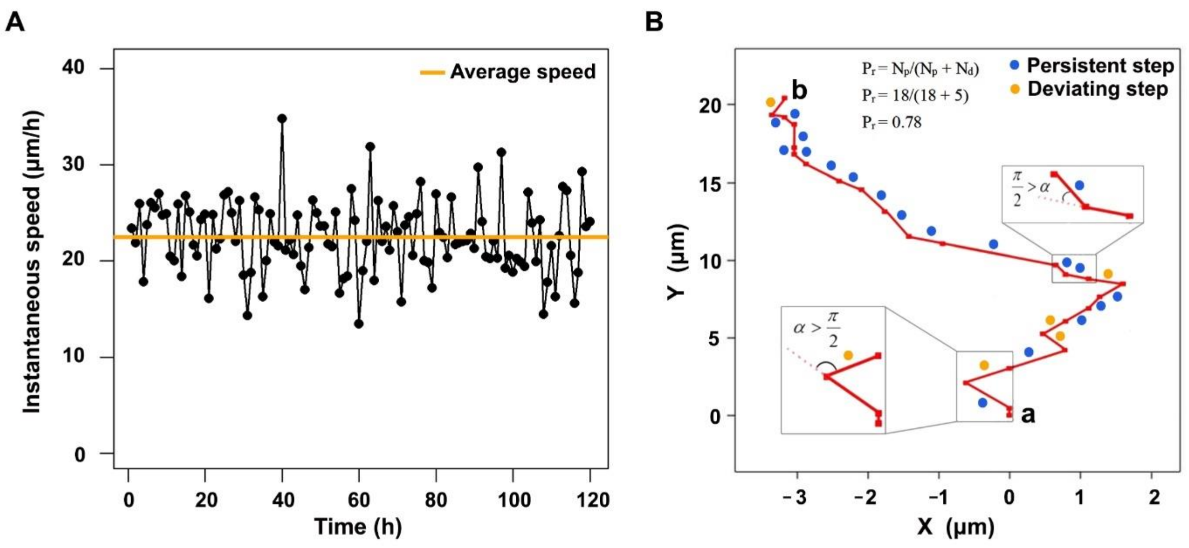


***Fig. S6. Cell's speed-persistence.*** *(A) 5-day speed profile. (B) Schematic chart explaining how the persistence ratio is computed for a cell that moved from a to b.*

iv) Spatial zones around lumina, referred to as well-nourished zones, are also modelled as squares of area W, with W > L, surrounding the lumina (Fig. S7A). When the position of an agent enters these zones, it has a higher probability to remain there than leaving to the poorly-nourished zone (w), that is the area among well-nourished zones mathematically described as:

$$w = F \setminus U_{i=1..Nv} W_i \qquad \text{(Equation S2)}$$

v) Cell division is a stochastic event. The probability of an agent to divide (Pd) is based on a proliferation score (PS) that depends on 3 additive factors: local cell density (D), the time since the division of the original agent (T) and proximity to a blood vessel (V). At each time point, these three factors are monitored for every cell in the simulated field. PS is computed as:

$$PS = D + T + V \qquad \text{(Equation S3)}$$

The local cell density (D) is a discrete index of the set  reflecting the number n=0,1,..,8 of occupied spots around the target simulated cell in Moore neighborhood regardless the location of the occupied spots (Fig. S7B). The higher the number n of the occupied spots in Moore neighborhood, the smaller is D.

D = 2   (if n = 0,1)

D = 1   (if n = 2,3)

D = 0   (if n = 4,5)

D = -1   (if n = 6,7)

D = -2   (if n = 8)

The proximity to a blood vessel (V) describes the proliferation advantage that a cell can get based on it is location. V is computed as:

$$V = B \qquad \text{(if } Z = 1\text{)} \qquad \text{(Equation S4)}$$

$$V = 0 \qquad \text{(if } Z = 0\text{)} \qquad \text{(Equation S5)}$$

where, Z refers to the location of the target cell in a poorly nourished zone w (Z = 0) or in a well-nourished zone $w^c$ (Z = 1), which is an advantage. This advantage is simulated as an additive bounce (B) that is equal to the time difference between the needed time for a cell to divide in a well-nourished zone ($d_{tw}$) and a poorly nourished zone ($d_t$). The proliferation score (PS) is converted to a high or low division probability ($P_d$) based on two additional elements; the density state (DS) and the doubling time, the duration of the entire cell cycle ($d_t$). The density state (DS) acts as a checkpoint assessing whether or not the simulated cell move forward with division.

$$B = d_t - d_{tw} \quad \text{(Equation S6)}$$

$$P_d = 0.001 \times DS \quad (\text{if } PS < d_t) \quad \text{(Equation S7)}$$

$$P_d = 0.999 \times DS \quad (\text{if } PS \geq d_t) \quad \text{(Equation S8)}$$

When a division takes place, two new daughter agents are generated. One agent keeps the position of the original agent. The second agent is placed stochastically based on a given speed and directionality and taking into consideration the available spots around the original agent before it divided. To achieve this, we assume that the second agent has moved from the location of the original agent by a distance computed based on the given speed. The direction of the movement is determined based on the given directionality. Putting together the computed distance and directionality, the center of the second agent can be any free point on the semicircle-circumference with a radius of the computed distance (Fig. S8). Points occupied by other cells are excluded before stochastically selecting the location of the second agent.

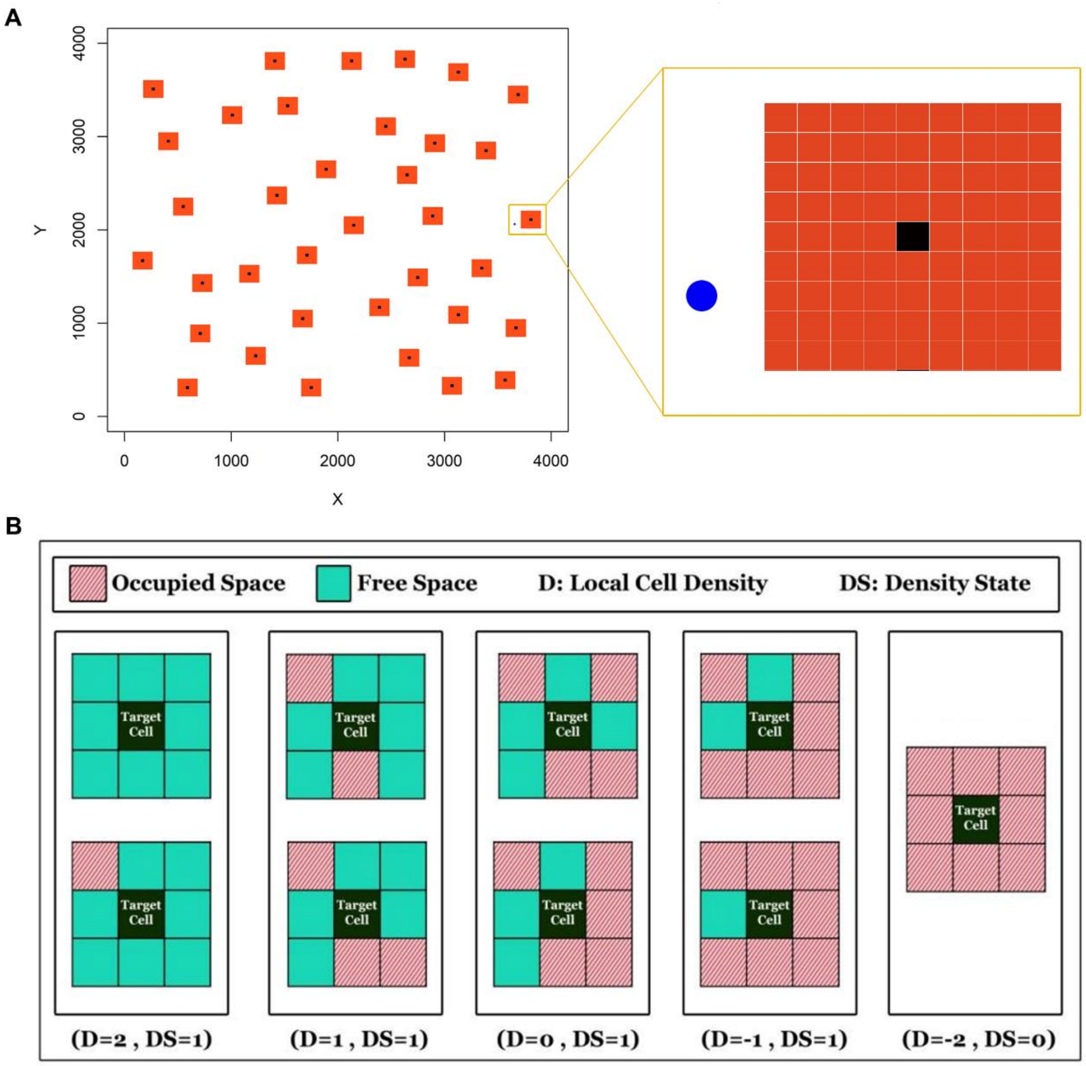

***Fig. S7. Simulation field and the local density.*** *(A) A 2D section showing the architecture of the simulated tumor field. Black squares represent the lumen of blood vessels. Red squares represent the well-nourished zones. The blue circle represents a single cancer cell. (B) Schema explaining the way of computing the local cell density and the density state.*

vi) The probability of undergoing cell division increases significantly when an optimal cell-cycle time is achieved. The duration of the cell cycle ($d_t$) is not only affected by the instantaneous presence in a well-nourished zone (Z = 1) but also by the cumulative time spent in a well-nourished zone. The first phase of the cell cycle is known as G1 phase. If the cell spends the entire G1 phase in a well-nourished zone then it gets an additive bounce (B). Thus, the cell can progress with the division regardless the Z value during the remaining cell cycle phases. To implement this bounce into the model we adjust the previous division probability:

$$P_d = 0.001 \times DS \quad (\text{if } t_{G1} > PS < d_t) \qquad \text{(Equation S9)}$$

$$P_d = 0.999 \times DS \quad (\text{if } t_{G1} \leq PS \geq d_t) \qquad \text{(Equation S10)}$$

where, $t_{G1}$ is the duration of G1 phase, $d_t$ *is* the duration of the entire cell cycle, PS is the proliferation score and DS the density state.

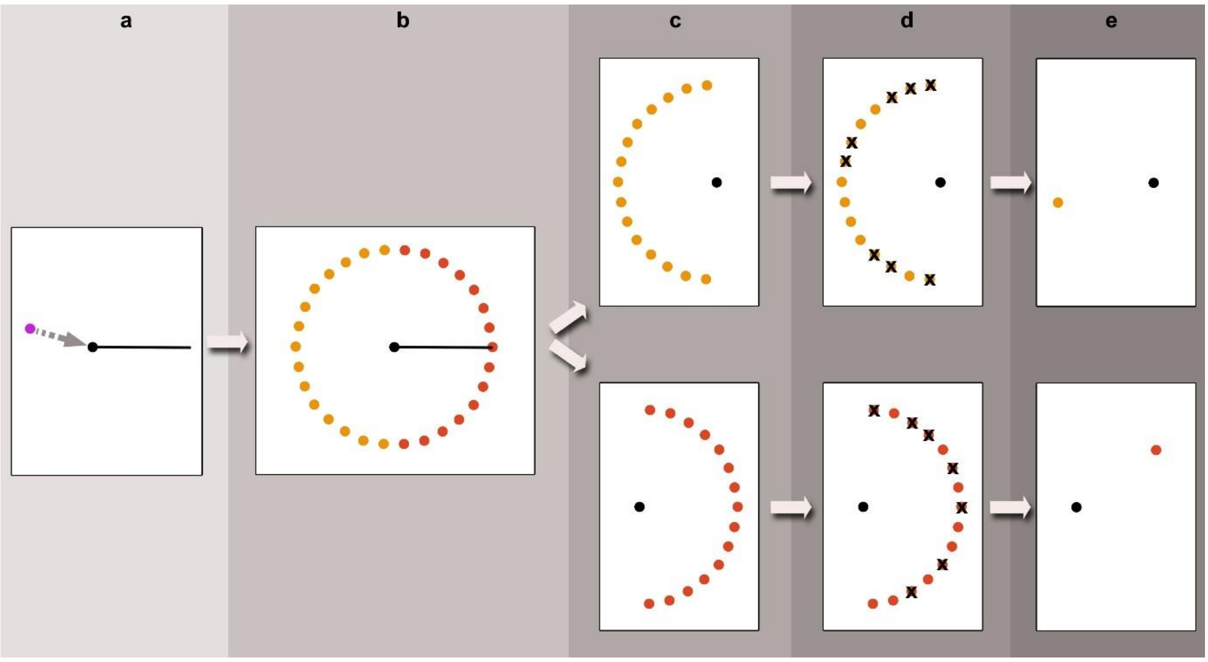


***Fig. S8. Simulating the position of the second daughter cell resulted from division of the original cell.*** *(A) Location of the original cell before (magenta) and after (black) division, the gray arrow shows the direction of the movement whereas the black line represents the distance between the second cell and the original cell. (B) All possible locations of the second cell. (C) Possible locations in case of forward movement (red dots) or backward movement (orange dots). (D) Excluding the spots occupied by other cells. (E) Stochastic selection of the location of the second cell in case of forward movement (red dot) or backward movement (orange dot)*

vii) Based on the lifespan length (T) of a cell, it can be classified as undergoing proliferation, senescent or necrotic. Cells cannot divide in the absence of available spots. Moreover, there is a low possibility for a cell with a sufficient proliferation score and availability of spots in the Moore neighborhood not to divide (Equation S10). On the other hand, under insufficient proliferation score, cells can still undergo division (Equation S9), as long as DS ≠ 0. If a cell is not undergoing cell division within a particular period of cell-cycles ($n_n$) then it is considered necrotic and removed from the field. Cells that did not divide within a given period of cell-cycles ($n_s$) are considered as senescent cells. The following conditions explain how the simulated cells are classified:

Undergoing proliferation if ($T \leq n_s \times d_t$ and $T \leq n_n \times d_t$)

*Senescent* if ($T > n_s \times d_t$ and $T < n_c \times d_t$)

*Necrotic* if ($T > n_n \times d_t$)

where, T is the time since the division of the original agent, $d_t$ is the duration of the entire cell cycle. Since in our model senescence is mainly dependent on the density state. Changes in the local cell density can shift a senescent cell back to a proliferating state or otherwise it becomes necrotic after a given period.

viii) Cells moving towards the lumina can intravasate with probability $P_i$. The lumen can either act as an absorbing barrier, if the cell succeeded entering the lumen, or as a reflecting barrier if the cell failed entering the lumen. If a cell failed entering the lumen, it stays in its previous location. On the other hand, intravasating cells are removed from the simulation field.

#### Model initialization and parameterization

Here we explain the choice of model parameters and initial conditions used in the simulations.

##### 2.1 Model initialization

For simplicity, we start the simulations with a single cell randomly placed at location ($X_0$,$Y_0$) within the 2D simulating field F. The coordinates of the first movement ($X_1$,$Y_1$) of both the tumor initiating cell and the newly generated cells are stochastically computed by generating a random number (rand) between 0 and 1 and using it according to the following equations:

$$X_1 = X_0 + [\text{cosine}(\text{rand} \times 2\pi) \times \text{distance}] \qquad \text{(Equation S11)}$$

$$Y_1 = Y_0 + [\text{sine}(\text{rand} \times 2\pi) \times \text{distance}] \qquad \text{(Equation S12)}$$

Within the simulation field, certain number of blood vessels (Nv) is stochastically distributed, assuming the spatial distribution of well-nourished zones surrounding blood vessels lumina is characterized by a minimum distance between any neighboring zones ($d_W$).

##### 2.2 Model parameters

A summary of all model parameters and values used is shown in Table1. Below we provide explanation of the choices made.

Table 1. A description of the parameters used in the proposed model.

| Parameter | Description | Value | Unit | Source |
|---|---|---|---|---|
| d | Cell diameter | 20 | µm | Empirical & estimated based on (1-4) |
| F | Simulation field area | 16 | $mm^2$ | Estimated based on ”d” |
| $L_i$ | Surface area of the lumen | 400 | $\mu m^2$ | Estimated based on ”d” |

| | | | | |
|---|---|---|---|---|
| $N_V$ | Number of blood vessels | 36 | Vessel count | Estimated based on (5) |
| Rrows | A number between 1 and 5 determining the number of red boxes in the four directions around the blood vessel. The total surface area of the well-nourished zones is computed as: $W = (N_V \times d^2 \times (2 \times Rrows+1)^2) - L$ | 4 | Rows around blood vessels | Estimated based on (6,7) |
| $d_t$ | Doubling time in poorly nourished zones. | 156 | Hours | Estimated based on (8) |
| $d_{tw}$ | Doubling time in well-nourished zones around blood vessels. | 108 | Hours | Empirical & estimated based on (8-10) |
| $n_n$ | Number of cell-cycle periods that leads to cell death if the division did not happen during that period. | 3 | Cell-cycle count | Estimated based on (11,12) |
| $n_s$ | Number of cell-cycle periods that leads to cell death if the division did not happen during that period. | 2 | Cell-cycle count | Scaled based on ” $n_n$” |
| LeavingRzone | Probability of leaving a well-nourished zone. (a value between 0 and 1)<br>For a cell to leave the well-nourished zone, it should get a random number that is equal to the "LeavingRzone " or less. | 0.5 | Dimensionless | Selected based on the outcome of sensitivity analysis |
| $P_i$ | Intravasation probability | 0.01 | Dimensionless | Estimated based on (13,14) |
| $I_{GF}$ | Intravasation Increment with Golgi fragmentation (A value between 0.5 and 1.2 multiplied by 33) | 33[0.5, 1.2] | Dimensionless | Empirical |
| G1 | Length of G1 phase | 60 | Hours | Estimated based on (8,15) |
| GFp | Golgi fragmentation frequency | 0, 14, 25, 50, 65, 75 & 100 | % | Empirical |
| $P_r$ | The persistence ratio value<br>(A value between 0 and 1) | Panel | Dimensionless | Empirical |
| Average Speed: minSpe | Minimum speed | Panel | um/h | Empirical |
| Average Speed: maxSpe | Maximum speed | Panel | um/h | Empirical |
| Average Speed: desired_meanSpe | Average speed | Panel | um/h | Empirical |
| Average Speed: desired_sdSpe | Speed standard deviation | Panel | um/h | Empirical |
| SimTime | Simulation time | 2160 | Hours | Inspired by literature (12) |
| TimeInterval | Time interval between steps | 1 | Hours | Scaled from fmpirical data |

- Cell diameter (d)

Clinical and experimental data of triple negative breast cancer reported a variety in cell size ranging from 10µm to 23µm in diameter (1-4). Based on the distribution of reported mean size of MDA-MB-231 in the previous studies and in-house microscopic observation **(**Fig. 9A**)**, we chose 20µm as the diameter of our simulated cells

- Simulation field area (F)

A tumor may contain many millions of cancer cells. Computational costs increase exponentially with the number of cells. For optimal use of available computational resources, we decided to simulate a cross-section (4mm × 4mm) of a tissue hosting a tumor. The maximum number of simulated cells at any time point is 40,000 cells.

- Surface are of the lumen ($L_i$)

For simplicity, we chose the lumen to have a square shape with a surface area of the square of the cell diameter.

- Number of blood vessels ($N_V$)

The $N_V$ was estimated based on the reported micro-vessel density in triple negative breast cancer patients (stages i and ii) (5).

- Rrows

This parameter is used to construct the well-nourished zone and it is estimated based on efficiency of the passive mode of molecular exchange through blood vessels. The efficiency was reported to high only over small distances (< 100um) (6,7).

- Doubling time in poorly nourished zones around blood vessels. *(*$d_t$*)*

The $d_t$ was set based on the reporte*d* doubling time of MDA-MB-231 cells in vivo immune-deficient NSG mice (NOD.Cg-*Prkdc*$^{scid}$ *Il2rg*$^{tm1Wjl}$/SzJ) which is about 6 days (11).

- Doubling time in well-nourished zones around blood vessels. *(*$d_{tw}$*)*

The $d_{tw}$ was estimated based on literature [8-10] in addition to in-house experimentation, where BT549 cells were cultured in RPMI medium containing 3%, 9% or 15% serum for 24 and 48h then cells were trypsinized and counted **(**Fig. S9B**)**.

- Number of needed arrested cell-cycle periods for necrosis to take place ($n_n$*)*

Taking together the estimation of doubling time in poorly nourished zones around blood vessels and the necrotic temporal conditions reported in (11,12), we estimated a period of 3 cell cycles without division for necrosis to take place.

- Number of needed arrested cell-cycle periods for senescence to take place ($n_s$*)*

Taking together the estimation of doubling time in poorly nourished zones around blood vessels and the estimation of ($n_n$*)*, we estimated a period of 2 cell cycles without division for senescence to take place.

- Probability of leaving *a* well-nourished zone *(*LeavingRzone*)*

The LeavingRzone was set to 0.5 meaning that there is 50% chance of accepting a movement of a cell in the well-nourished zone to go out. Sensitivity analysis showed that changing the value of the LeavingRzone does not change the trend of the results.

- Intravasation probability ($P_i$)

The ($P_i$) was estimated based on literature (13,14). Although the number of cells increases with the simulation time, the ($P_i$) was kept constant during the simulation. Because of the minor change of intravasation rate with increased tumor size in breast cancer (16,17).

- Intravasation Increment with Golgi fragmentation ($I_{GF}$)

Our experimentation showed that Golgi fragmentation increases the intravasation rate 33-fold (Fig. S9C). For a better mimicking of the empirical data, instead of using a single value, the $I_{GF}$ stochastically can take a value within a range. This value will be used to adjust the intravasation rate:

$$\text{Intravasation Rate}_{\text{(with Golgi fragmentation)}} = P_i \quad I_{GF} \times 33 \qquad \text{(S11)}$$

- Length of G1 phase (G1)

The Length of G1 phase was reported to be (18–24 h) (15), by scaling it based on the total cell cycle length in vivo [8], we chose 60h as a good value for G1.

- Golgi fragmentation frequency (GFp)

The GFp is a panel of values 0, 14, 25, 50, 65, 75 & 100. Only 14 and 65 were estimated empirically. However, the other values were selected for comparison purposes.

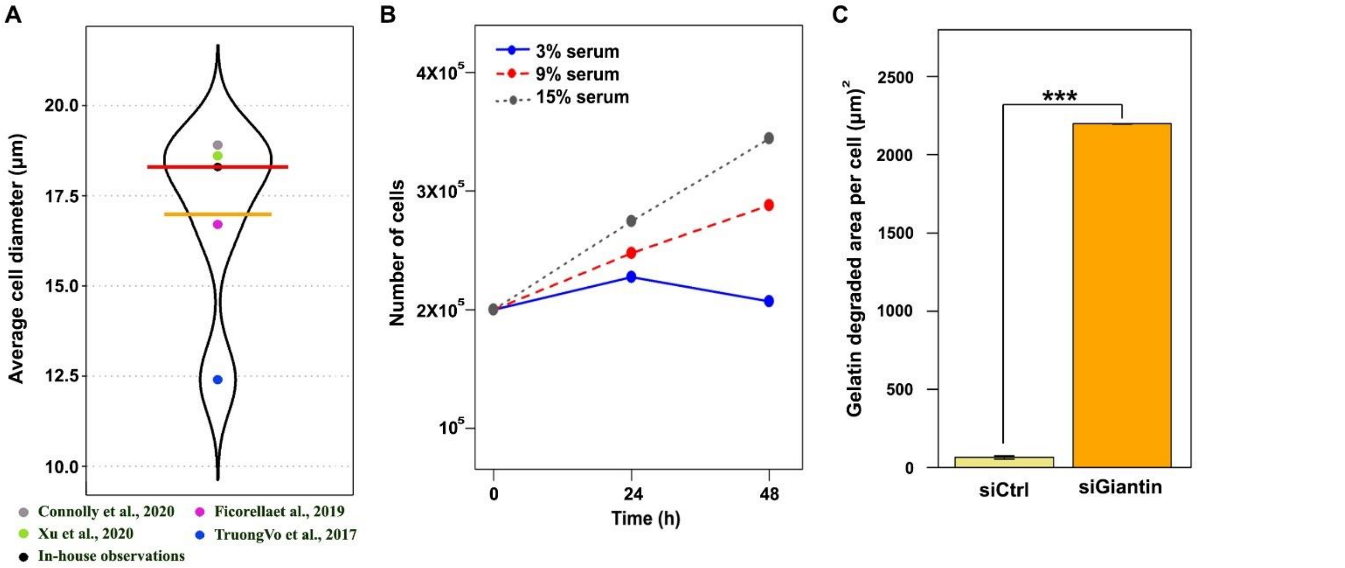


***Fig. S9. Experimental observations of BT548 and MDA-MB-231 cells.*** *(A) Violin plot showing of reported mean diameter of MDA-MB-231 in the several studies and in-house microscopic observation. Median cell diameter is shown in red whereas the mean is shown in orange. (B) line plot showing the number of BT549 cells* cultured in RPMI medium containing 3%, 9% or 15% serum *at three different time points. (C) Quantification of the gelatin-degraded area. Bars represent the median value. Error bars represent quantile-based coefficient of variation relative to control cells by Kruskal-Wallis chi-squared test (**** $p < 0.001$*), n=13 microscopic fields from 2 independent experiments.*

- The persistence ratio $P_r$ and average speed

In order to mimic the movement of both MDA-MB-231 and BT549 cells, we grow them in 2D cultures by seeding $20 \times 10^3$ mother cells and $5 \times 10^3$ cells stably expressing fluorescent proteins that localize to the nucleus. Cells with fluorescent nuclei were tracked analyzed using our R package cellmigRation (18). In total, 1398 MDA-MB-231 cells and 2850 BT549 cells were analyzed. Based on the violin plots of the speed and persistence ratio of cells (Fig. S10A&B), a set of 10 persistence values and 13 speeds were sampled from the distribution of empirical values to be used as input for the modeling with 130 speed-persistence combinations generated by the Cartesian product of the persistence and speed sets (Fig. S10C). The experimental data showed a significant moderated correlation between speed and persistence ratio ($r_{BT549} = 0.44$, $r_{MDA\_MB\_231} = 0.47$, p-value $< 0.001$) (Fig. S10D). Based on our experimental observation, we noticed that cells move either in the same direction as the previous movement (Fig. S10E) within bilateral 90 degrees (red semi-circle) or in opposite direction to the previous movement within bilateral angles between 90 and 180 degrees (orange semi-circle).

- Simulation time (SimTime)

The number of cells increases very fast with time, dramatically increasing the needed computational costs. Thus, we set the simulation time to 90 days inspired by literature (12).

- Time interval between steps (Timeinterval)

In our experimentation we used a time interval of 10min. Due to the computational cost of applying short time intervals, we decided to set the time interval to 1hour.

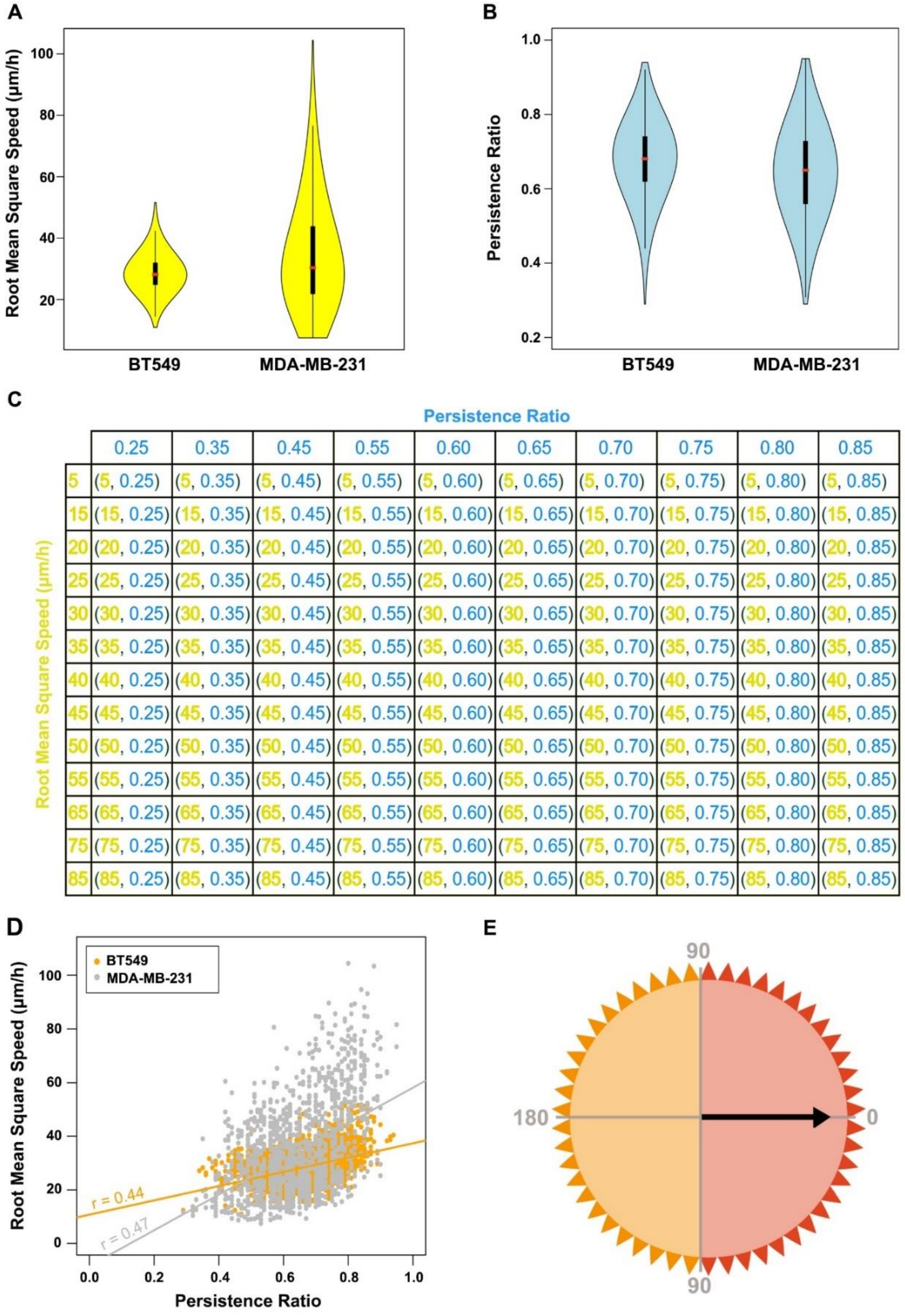


| | 0.25 | 0.35 | 0.45 | 0.55 | 0.60 | 0.65 | 0.70 | 0.75 | 0.80 | 0.85 |
|---|---|---|---|---|---|---|---|---|---|---|
| 5 | (5, 0.25) | (5, 0.35) | (5, 0.45) | (5, 0.55) | (5, 0.60) | (5, 0.65) | (5, 0.70) | (5, 0.75) | (5, 0.80) | (5, 0.85) |
| 15 | (15, 0.25) | (15, 0.35) | (15, 0.45) | (15, 0.55) | (15, 0.60) | (15, 0.65) | (15, 0.70) | (15, 0.75) | (15, 0.80) | (15, 0.85) |
| 20 | (20, 0.25) | (20, 0.35) | (20, 0.45) | (20, 0.55) | (20, 0.60) | (20, 0.65) | (20, 0.70) | (20, 0.75) | (20, 0.80) | (20, 0.85) |
| 25 | (25, 0.25) | (25, 0.35) | (25, 0.45) | (25, 0.55) | (25, 0.60) | (25, 0.65) | (25, 0.70) | (25, 0.75) | (25, 0.80) | (25, 0.85) |
| 30 | (30, 0.25) | (30, 0.35) | (30, 0.45) | (30, 0.55) | (30, 0.60) | (30, 0.65) | (30, 0.70) | (30, 0.75) | (30, 0.80) | (30, 0.85) |
| 35 | (35, 0.25) | (35, 0.35) | (35, 0.45) | (35, 0.55) | (35, 0.60) | (35, 0.65) | (35, 0.70) | (35, 0.75) | (35, 0.80) | (35, 0.85) |
| 40 | (40, 0.25) | (40, 0.35) | (40, 0.45) | (40, 0.55) | (40, 0.60) | (40, 0.65) | (40, 0.70) | (40, 0.75) | (40, 0.80) | (40, 0.85) |
| 45 | (45, 0.25) | (45, 0.35) | (45, 0.45) | (45, 0.55) | (45, 0.60) | (45, 0.65) | (45, 0.70) | (45, 0.75) | (45, 0.80) | (45, 0.85) |
| 50 | (50, 0.25) | (50, 0.35) | (50, 0.45) | (50, 0.55) | (50, 0.60) | (50, 0.65) | (50, 0.70) | (50, 0.75) | (50, 0.80) | (50, 0.85) |
| 55 | (55, 0.25) | (55, 0.35) | (55, 0.45) | (55, 0.55) | (55, 0.60) | (55, 0.65) | (55, 0.70) | (55, 0.75) | (55, 0.80) | (55, 0.85) |
| 65 | (65, 0.25) | (65, 0.35) | (65, 0.45) | (65, 0.55) | (65, 0.60) | (65, 0.65) | (65, 0.70) | (65, 0.75) | (65, 0.80) | (65, 0.85) |
| 75 | (75, 0.25) | (75, 0.35) | (75, 0.45) | (75, 0.55) | (75, 0.60) | (75, 0.65) | (75, 0.70) | (75, 0.75) | (75, 0.80) | (75, 0.85) |
| 85 | (85, 0.25) | (85, 0.35) | (85, 0.45) | (85, 0.55) | (85, 0.60) | (85, 0.65) | (85, 0.70) | (85, 0.75) | (85, 0.80) | (85, 0.85) |

***Fig. S10. Experimental observations of Bt548 and MDA-MB-231 cells.*** *(A) Violin plot showing the distribution of the values of root mean square speed for BT549 and MDA_MB_231 cells. (B) Violin plot showing the distribution of the values of persistence ratio for BT549 and MDA_MB_231 cells. (C) Cartesian product of the persistence ratio and the root mean square speeds used for the modeling. (D) A scatterplot showing the relationship between the persistence ratio and the root mean square speeds of BT549 cells (orange) and MDA_MB_231 cells (gray). (E) Movement options of simulated cells. Black arrow represents the previous movement. Red triangles represent the possible locations of a forward movement whereas orange triangles represent the possible locations of a backward movement.*

#### 3. Simulation set-up and additional computational details

In order to mimic the behavior of breast cancer cells computationally, we selected simulation scenarios based on experimental observations. We constructed the proposed model to imitate as much as feasible the pathophysiological traits of a low-grade triple-negative breast cancer based on in-house experimentation and inspired by literature. However, the parametric structure of our model makes it applicable to different tumors simply by setting optimal values for a panel of parameters, such as the number of blood vessels, simulation time, cell speed and Golgi fragmentation frequency. In table 2, we demonstrate the parameters used in the proposed model with their corresponding computational name.

In our model, the ability of a cell to move, divide, intravasate and leave a well-nourished zone is a stochastic event. Such stochasticity is achieved by generating a random number ($R \in [0,1]$) per cell per process per iteration. A process is triggered when the random number is equal to or less than the corresponding parameter value. For example, to compute the direction of movement, a random number between 0 and 1 is generated. In case the random number is equal or greater than the $P_r$, then the cell will move in opposite direction to the previous movement. However, if the random number is less than the $P_r$, then the cell will move in same direction as the previous movement. To achieve this, we assume that the new location is a point of a set of points on the circle circumference with a radius of a particular distance computed from the speed profile (Fig. S6A). To get these points, first a series of trigonometry equations (Fig. S11A) are used to compute the coordinates ($x_3$,$y_3$) of the outermost point of the radius ($p_3$) that forms a right angle with the distance of the previous movement ($D_1$). Then, the coordinates of the points on the circle circumference are computed using the following equations:

$$X_i = x_2 + \text{cosine}(i \times \pi / 180) \times (x_3 - x_2) - \text{sine}(i \times \pi / 180) \times (y_3 - y_2) \quad \text{(Equation S14)}$$

$$Y_i = y_2 + \text{cosine}(i \times \pi / 180) \times (y_3 - y_2) - \text{sine}(i \times \pi / 180) \times (x_3 - x_2) \quad \text{(Equation S15)}$$

Where i is a set of angles. In case of forward migration, the set contains 44 angles between 184° and 356° to keep the direction of the movement within bilateral 90 degrees whereas in backward migration the set contains 46 angles between -180° and -360° (Fig. S11B). Finally, the grid-based spatial reference is used to select one point stochastically among the available points that are not occupied by other cells (Fig. S11C). In case all the points are occupied, the cell will stay in place.

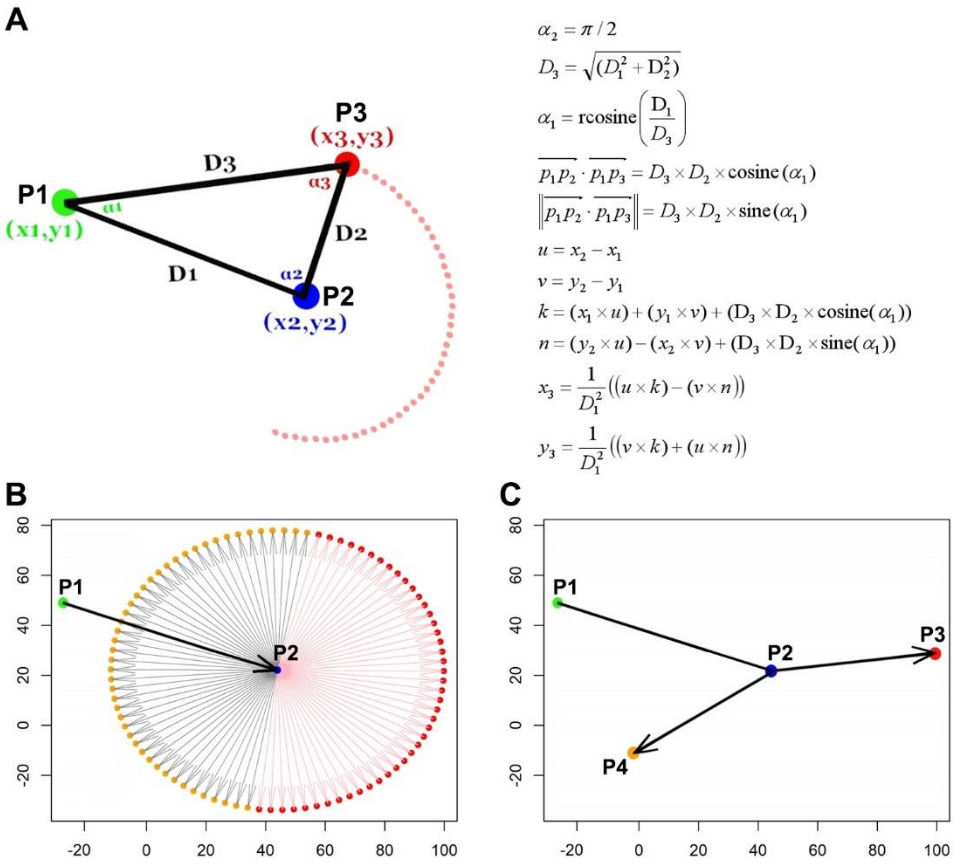


*__Fig. S11. Simulating cell movements.__ (A) A schema with a panel of mathematical equations explaining how the X and Y coordinates of the new movement are computed. (B) Possible locations in case of forward movement (red dots) or backward movement (orange points). (C) A stochastic selection of a forward movement from P2 to P3 and a backward movement from P2 to P4.*

For navigation purposes, a grid-based spatial reference system was embedded on top of the 2D simulation field. The number of grid squares in each direction of the computational grid is determined by dividing the length of simulation field side by the diameter of the simulated cancer cell generating a grid of 40,000 squares. Each grid square is 20µm × 20µm. This lattice grid simply helps determining the cell local density around a target cell using the Moore neighborhood method *(*Fig. S12A*)*. Since a small spatial field of the tumor is simulated, periodic boundary conditions were imposed to minimize the edge effect and avoid losing cells (19). For example, if a cell migrates past the left edge, it will then appear at the right edge (Fig. S12B). Lattice squares are numbered from 1 to 40,000, where 1 is located at the lower left corner whereas 40,000 is located at the upper right corner (Fig. S12C). The lattice square numbering increases from left to right and bottom to top.

To evaluate the distribution of the simulated cancer cells, the uniformity index (UI) is computed as the average of 16 sectional uniformity indices ($UI_i$) representing the 16 sections of the simulated field (Fig. S12D).

$$UI = \frac{\sum_{i=1}^{16} UI_i}{16} \qquad \text{(Equation S16)}$$

Each sectional uniformity index ($UI_i$) is computed as one minus coefficient of variation (CV) multiplied by hundred. The CV represents the ratio of the standard deviation to the mean of the occupied grid squares within the section.

$$UI_i = (1 - CV) \times 100$$ ( Equation S17)

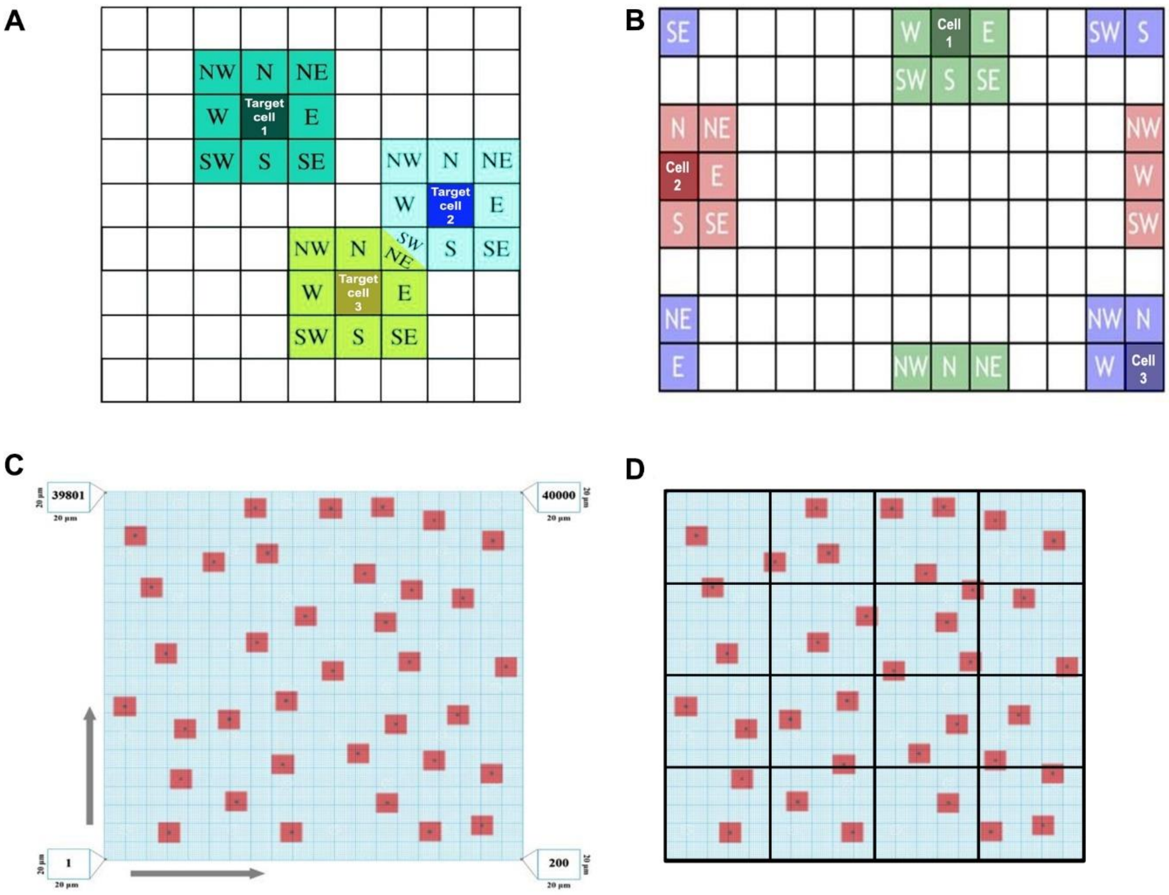


***Fig. S12. The simulation field.*** (A) *Moore neighborhood. (B) Moore neighborhood in periodic boundary conditions. (C) The grid-based spatial reference system embedded on top of the 2D simulation field, arrows show the increase in the lattice square numbering.*

All the modeling simulations were implemented using the high-performance computing at Oslo University (Abel computer cluster). The source code of the proposed model is available at https://github.com/ocbe-uio/Cancer_simulator.

Table 2. Parameters used in the proposed model with their corresponding computational name.

| Parameter name | Corresponding computational name |
|---|---|
| d | _ |
| F | _ |
| $L_i$ | _ |
| $N_V$ | BloVesN |
| Rrows | Rrows |
| $d_t$ | DT |

| | |
|---|---|
| $d_{tw}$ | DTinR |
| $n_n$ | PeriodForDeath |
| $n_s$ | PeriodForSenescence |
| LeavingRzone | LeavingRzone |
| Intravasation probability ($P_i$) | IntravasationProb |
| Intravasation Increment with Golgi fragmentation ($I_{GF}$) | IntraVas_inc_By_GF |
| G1 | G1 |
| GFp | Golgi_Fra_Frequency |
| $P_r$ | PER |
| minSpe | minSpe |
| maxSpe | maxSpe |
| desired_meanSpe | desired_meanSpe |
| desired_sdSpe | desired_sdSpe |
| SimTime | SimTime |
| TimeInterval | TimeInterval |
